# Shape variability of the central sulcus in the developing brain: a longitudinal descriptive and predictive study in preterm infants

**DOI:** 10.1101/2021.12.15.472770

**Authors:** H de Vareilles, D Rivière, Z Sun, C Fischer, F Leroy, S Neumane, N Stopar, R Eijsermans, M Ballu, ML Tataranno, MJNL Benders, JF Mangin, J Dubois

**Affiliations:** Université Paris-Saclay, NeuroSpin-BAOBAB, CEA, Gif-sur-Yvette, France; Université Paris-Saclay, NeuroSpin-UNICOG, Inserm, CEA, Gif-sur-Yvette, France; Université de Paris, NeuroDiderot, Inserm, Paris, France; Université Paris-Saclay, NeuroSpin-UNIACT, CEA, Gif-sur-Yvette, France; Utrecht University, University Medical Center Utrecht, Department of Neonatology, Utrecht, the Netherlands; Department of Pure Mathematics and Mathematical Statistics, University of Cambridge, Cambridge, United Kingdom

**Keywords:** Brain morphological development, cortical folding, hemispheric asymmetries, prematurity, outcome prediction, lateralization, fine motor outcome, manual dexterity.

## Abstract

Despite growing evidence of links between sulcation and function in the adult brain, the folding dynamics, occurring mostly before normal-term-birth, is vastly unknown. Looking into the development of cortical sulci in babies can give us keys to address fundamental questions: what is the sulcal shape variability in the developing brain? When are the shape features encoded? How are these morphological parameters related to further functional development?

In this study, we aimed to investigate the shape variability of the developing central sulcus, which is the frontier between the primary somatosensory and motor cortices. We studied a cohort of 71 extremely preterm infants scanned twice using MRI – once around 30 weeks post-menstrual age (w PMA) and once at term-equivalent age, around 40w PMA –, in order to quantify the sulcus’s shape variability using manifold learning, regardless of age-group or hemisphere. We then used these shape descriptors to evaluate the sulcus’s variability at both ages and to assess hemispheric and age-group specificities. This led us to propose a description of ten shape features capturing the variability in the central sulcus of preterm infants. Our results suggested that most of these features (8/10) are encoded as early as 30w PMA. We unprecedentedly observed hemispheric asymmetries at both ages, and the one captured at term-equivalent age seems to correspond with the asymmetry pattern previously reported in adults. We further trained classifiers in order to explore the predictive value of these shape features on manual performance at 5 years of age (handedness and fine motor outcome). The central sulcus’s shape alone showed a limited but relevant predictive capacity in both cases. The study of sulcal shape features during early neurodevelopment may participate to a better comprehension of the complex links between morphological and functional organization of the developing brain.

**Highlights:**

- Shape features can be isolated to describe quantitatively the development of the central sulcus.
- Most shape characteristics of the central sulcus are already encoded at 30 weeks of post-menstrual age (w PMA) in preterm newborns.
- The central sulcus shows subtle hemispheric asymmetries as soon as 30w PMA.
- The early shape of the central sulcus can help predicting handedness and fine motor outcome at 5 years of age.

## Introduction

The folding of the brain is highly complex: it shows similar patterns within all humans, yet it is unique for each individual. Even though its general aspect has been studied and described centuries ago (Cunningham, 1892), the inter-individual variability in sulcation has historically been eluded due to its complexity. More recently, this topic has made its appearance in literature, but in the vast majority of cases, this variability was addressed as a problem for group studies and the purpose was to dim it out (Fox et al., 1985). In the last few decades, we have witnessed the emergence of studies focusing on the description of sulcal pattern variability (Ono et al., 1990; Welker, 1990; Mangin et al., 2015). This new trend can be explained by multiple advances, among which the ones in magnetic resonance imaging (MRI), allowing non-invasive imaging of the brain of both pathological and healthy subjects, and the advances in machine learning, which enables the scientific community to address complex questions which used to be out of the reach of human comprehension.

Different observations have suggested that the shape of sulci can be linked to function, both in healthy and pathological cases. The most obvious examples reside in pathologies with strong sulcal malformation, such as lissencephaly (resulting in a flat brain with no sulci) or polymicrogyria (resulting in supernumerary unusually small convolutions). In less obviously sulcation-linked pathologies, a vast quantity of studies have investigated basic shape markers such as sulcal length or depth, easily quantifiable (Cachia et al., 2008). Subtle alterations in folding have also been highlighted based on complex shape variability, embedded in the pattern of folds in some neurological and psychiatric diseases, such as epilepsy or schizophrenia (Plaze et al., 2011; Mellerio et al., 2015). Variations in sulcal patterns linked to function have also been observed in healthy populations. For instance, the sulcal pattern of the anterior cingulate cortex has been reported to predict cognitive control performance across lifespan, and the favorable pattern for cognitive control has been reported to be the opposite in the monolingual and bilingual populations (Cachia et al., 2017).

These different elements give us keys to address fundamental questions about sulcation: can we differentiate a healthy variability in the folding pattern of the brain versus a pathological one? What are the functional implications of this variability?

To explore these questions, we focused this study on a single sulcus, to characterize its folding pattern during development and to look into its relationships with clinical and functional outcomes. The central sulcus was an excellent candidate for this purpose. First of all, it is a primary fold that appears as early as 20 weeks of GA during development (Chi et al., 1977), with potential long-lasting developmental alterations in cases of early disturbance during gestation or perinatal period (e.g. early brain injury, preterm birth). Moreover, it appears to be very stable across individuals, which makes it easy to identify even in very young preterm infants. Secondly, the central sulcus represents the anatomical border between two highly specialized cortical regions: the primary motor and somatosensory cortices. A detailed functional organization has been described into both areas (i.e. somatotopic motor and sensory maps) in adults (Penfield and Rasmussen, 1950; Germann et al., 2019), such that we can identify precise correspondent subregions of the central sulcus related to specific body parts. Some studies have already shown strong relationships between the central sulcus shape and some functional characteristics in adults (Sun et al., 2012; Sun et al., 2016; Germann et al., 2019; Mangin et al., 2019). For example, the location and shape of the upper knob of the central sulcus, commonly referred to as the “hand-knob”, has been related to the functional hand region (Yousry et al., 1997; Sastre-Janer et al., 1998; Sun et al., 2016), with differences observed between left and right-handers (Sun et al., 2012).

It should be noted that during the third trimester of pregnancy, the brain undergoes drastic changes, on both the macroscopic and microscopic scales. Many concurrent events occur, key to the development of the brain structure and function, among which the development of sulcation, with the brain progressively switching from the state of slightly dimpled to a well-folded state (Dubois et al., 2019; Kostović et al., 2019). At full-term birth, the brain of a baby roughly resembles that of an adult in terms of complexity and advancement of folding, and even though the shape of the sulci keeps on evolving throughout infancy and lifetime, the changes remain subtle compared to the ones observed during late gestation. Therefore, since sulcation is well developed at full-term birth, one can wonder when the general sulcal patterns emerge. In order to look into this question in normal development, it would seem sensible to study sulcal development in healthy fetuses. Yet, longitudinal studies of sulcation on healthy fetuses are currently highly challenging because of the difficulties encountered in acquisition and processing of fetal images at a sufficient resolution to study subtle shape variations in folding. Moreover, it has been reported that birth affects the shape and folding of the brain (Lefèvre et al., 2016), which makes it even more challenging to consider longitudinal studies comparing pre- and post-natal images. Therefore, an interesting alternative is to study the shape of sulci in preterm neonates before full-term equivalent age, even though their sulcation is somehow altered by prematurity (Bouyssi-Kobar et al., 2016; Dubois et al., 2019).

Studying the sulcation at different time-points in infants born preterm allows us to investigate longitudinally the dynamics of early sulcation, and to explore the effects of early brain adversities on these mechanisms. Very preterm infants are at risk of a wide variety of neurodevelopmental disorders, including motor disabilities and cerebral palsy (Korzeniewski et al., 2018; Pascal et al., 2018). The sensorimotor cortex is already well developed and connected at full-term birth, while it is still vastly developing in the case of very preterm births. Extreme prematurity therefore exposes the sensorimotor cortex to adversities when it is most vulnerable, and still growing, which could have parallel effects on function and sulcation (Yamada et al., 2016).

In a previous study, we assessed the relationship between early cortical changes and neurodevelopmental outcome in extremely preterm infants, using basic shape markers (sulcal surface area and mean depth) (Kersbergen et al., 2016). This study reported some links between sulcation and outcome, particularly between the inferior frontal sulcus and receptive language score at 2 years of age. But no links were captured between the central sulcus and the motor outcome, suggesting that basic shape markers might not be sufficient to capture relevant folding variability for the motor outcome. Regarding the sensorimotor region, early developmental disturbances have long-term consequences on brain morphology. For example, congenital one-hander adult participants showed a significantly flatter central sulcus in the contralateral hemisphere compared to both individuals with acquired upper-limb amputation and control subjects (Sun et al., 2017). This suggested that the mature sulcus’s shape is influenced by early sensorimotor activity and by the development of brain connectivity. This led us to the hypothesis that some specific folding variability observed in the central sulcus of preterm infants, resulting from premature birth and strong differences in *ex utero* perceptions, movements and environment, might be characteristic of deviations in normal brain development impacting their functional outcome. Therefore, quantifying the links between the early central sulcus’s shape and the infant’s motor outcome at a later age could allow both a better understanding of the early functional implications of shape variability and the identification of early informative biomarkers for earlier diagnosis and accurate follow-up of these at-risk infants.

In this study, our aim was therefore to capture the early central sulcus’s shape variability from a longitudinal cohort of extremely preterm infants, and to link it to their fine motor outcome at 5 years of age. In order to do so, we used longitudinally acquired MRI data at two specific ages: one at around 30 weeks of post-menstrual age (w PMA), when the central sulcus is already present but at an early stage of development, and one around term-equivalent age (TEA, at around 40w PMA), when the central sulcus is well developed. We then characterized the shape variability of the central sulcus in this cohort, explored the relationships between its shape at 30w PMA and at 40w PMA, and investigated the shape predictive capacity on the motor development of children evaluated at around 5 years of age.

## Material and methods

### Subjects and preprocessing of brain images

### Cohort and collection of clinical and outcome data

The study was carried out on a retrospective longitudinal cohort of extremely preterm infants (Kersbergen et al., 2016). Permission from the medical ethics review committee was obtained for this study. A total of 71 subjects (51% males) from the Wilhelmina’s Children Hospital, University Medical Center, Utrecht, born between 24 and 28 weeks of gestational age were included.

In agreement with our previous study (Kersbergen et al., 2016), the perinatal clinical factors of interest retained for the analyses were the gestational age at birth (GA), the birth-weight z-score (BWZ) computed according to the Dutch Perinatal registry reference data (Visser et al., 2009), and the presence of an intra-ventricular hemorrhage (IVH, dichotomized between absent or mild (no IVH or IVH of grades 1 - 2 according to the Papile scale) and severe (grades 3 - 4)). We also recorded the presence of broncho-pulmonar dysplasia (BPD).

Around five years of age (age range: 4 years and 6 months – 6 years and 7 months, mean: 5 years and 9 months ± 4 months), the children’s handedness was assessed by a trained pediatric physical therapist who asked them first either to write their name or draw a picture, and secondly to pick up a coin placed at the center of the table. If the two tasks were done using the same hand, the hand preference was set to either left or right, otherwise it was reported as mixed. The parents were then asked if this hand preference matched that observed at home while writing or drawing. If the parents disagreed, the hand preference was set to mixed. At the same age, 66 out of the 71 children underwent the Movement Assessment Battery for Children, 2^nd^ edition, Dutch version (MABC-2- NL). The scores were corrected for age at evaluation. The assessment included tests for balance, aiming/catching and manual dexterity. See Sup. Info., Table 1 for the description of these clinical factors and outcome scores.

**Table 1.**
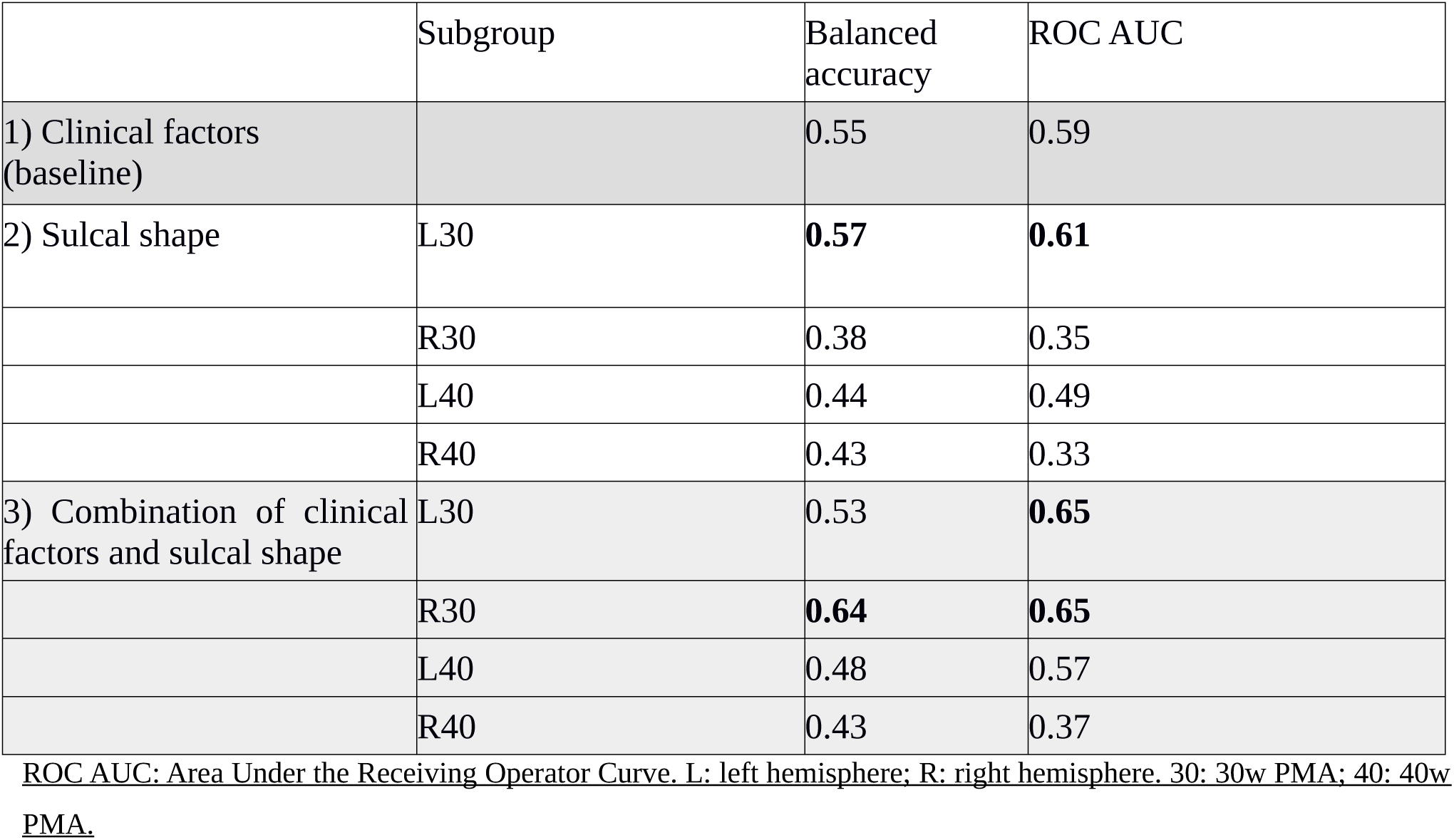
Cross-validated scores for handedness classification. Scores obtained using 1) clinical factors; 2) sulcal shape features (positions corrected for PMA on the 10 dimensions), for each age and side subgroup; and 3) combination of both (clinical factors and sulcal shape features), for the handedness discrimination at 5 years of age. The bold values are the ones equal or superior to the baseline, i. e. the values obtained for the clinical factors alone.

### MRI data acquisition

The infants were scanned twice soon after birth: once around 30w PMA (28.7 − 32.7w, mean: 30.7 ± 0.9w) and again around term equivalent age (TEA) (40.0 − 42.7w, mean: 41.2 ± 0.6w). For each scan, MR imaging was performed on a 3-Tesla MR system (Achieva, Philips Medical Systems, Best, The Netherlands). The protocol included T2-weighted imaging with a turbo-spin echo sequence in the coronal plane (at early MRI: repetition time (TR) 10.085 ms; echo time (TE) 120 ms; slice thickness 2 mm, in-plane spatial resolution 0.35 × 0.35 mm; at TEA: TR 4847 ms; TE 150 ms; slice thickness 1.2 mm, in-plane spatial resolution 0.35 × 0.35 mm).

### Preprocessing of brain images

The preprocessing of MRI images was performed following the methodology already described in a previous study (Kersbergen et al., 2016): after generating a brain mask, T2-weighted images were segmented into three classes (grey matter, unmyelinated white matter and cerebrospinal fluid) using supervised voxel classification (Moeskops et al., 2015). By adapting the BabySeg and Morphologist anatomical pipelines of the BrainVISA software (BrainVISA suite, https://brainvisa.info), these segmentations allowed a reconstruction of the inner cortical surfaces of both hemispheres, and the extraction of objects depicting the sulci. A summary of the pipeline and the resulting 3D visualizations of the left hemisphere of a subject are presented in Figure 1.

**Figure 1.**
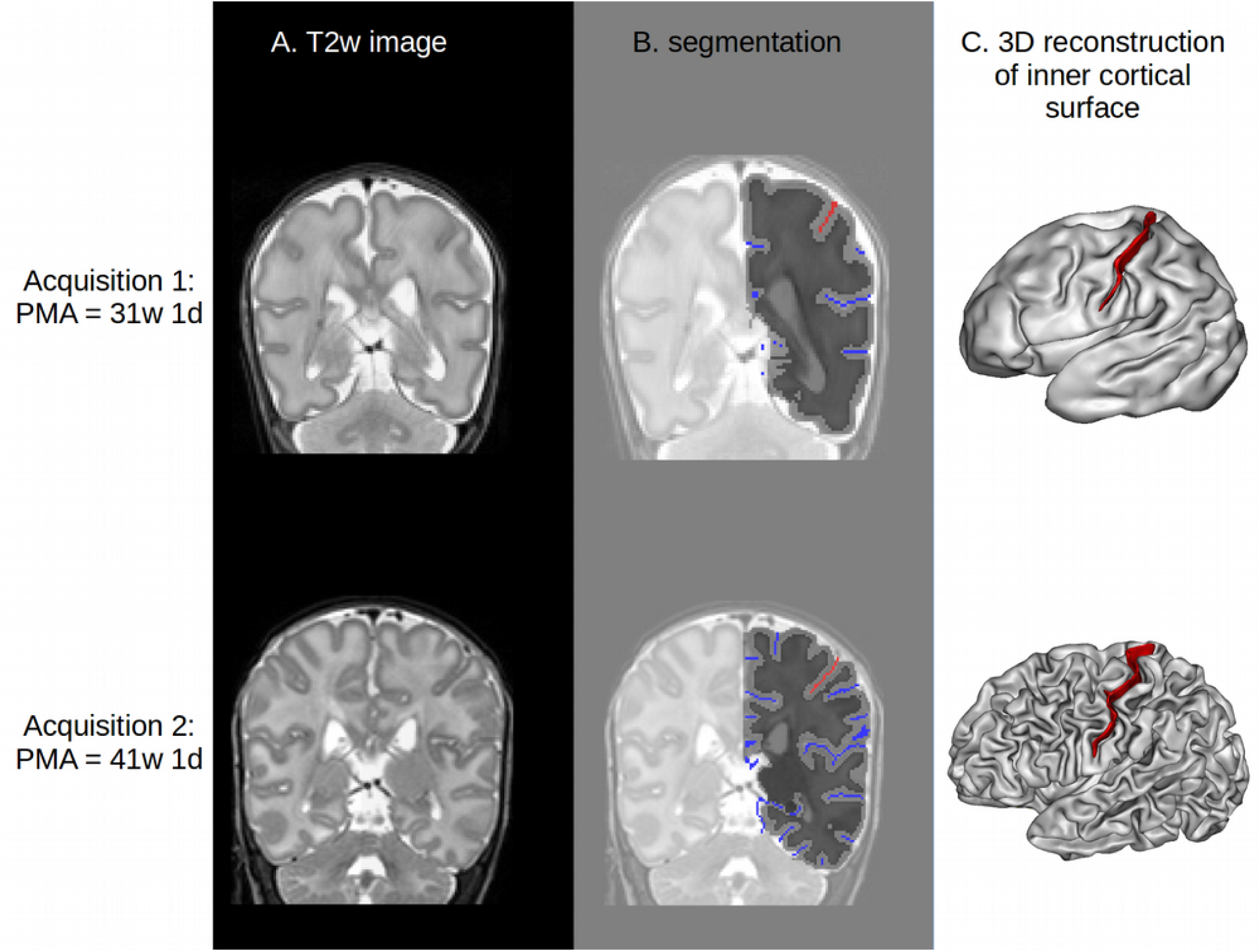
Summary of pipeline for sulcal extraction. A: sample T2-weighted coronal MRI. B: corresponding segmentation of the left hemisphere, showing the boundary between white (dark grey) and grey (light grey) matter. The sulci are represented in blue, except for the central sulcus, in red. C: 3D representation of the reconstructed surface of the white matter. Red ribbon: central sulcus.

### Characterization of the central sulcus’s shape

### Purpose of shape characterization

The purpose of shape characterization on the central sulcus is first to identify the shape features covering the main variability across infants (descriptive approach), and second to quantify the individual specificities relative to these shape features (comparative approach). The method used for shape characterization is inspired by a method already deployed on adults (Sun et al., 2016). A previous study in this cohort (Kersbergen et al., 2016) already addressed the link between sulcation (including the central sulcus) and clinical outcome, but using simpler metrics to quantify sulcal shape, such as surface area or depth. The added value of the current study is to improve the accuracy of sulcal shape assessment.

### Sulci extraction

First, all the cortical folds were extracted using the Morphologist toolbox of the BrainVISA software in the two serial images of each subject. Each fold was materialized by the skeleton of the volume of its cerebrospinal fluid, namely a set of voxels sampling a surface equidistant to the walls of the two adjacent gyri (Mangin et al., 1995). Each sulcus of the anatomical nomenclature corresponds to a set of the elementary folds extracted during this initial stage. The sulci labels were then identified automatically using a Bayesian pattern recognition strategy (Perrot et al., 2011). Finally, the central sulci labels were manually checked and corrected when necessary by one of the authors (HV). In order to dim out the important variability in cerebral size and shape, particularly between the two sessions of investigation (early at ∼30w PMA, vs TEA at ∼40w PMA), the obtained central sulci representations were transformed to the Talairach space using an affine transformation. The right sulci were finally mirrored relative to the interhemispheric plane in order to facilitate the comparison with the left ones.

### Sulci co-registration

The previous step led to the extraction of 284 central sulci (71 subjects x 2 hemispheres x 2 acquisitions). The dissimilarity between any two given sulci was then computed in the following way: sulcus A was registered to sulcus B using a rigid transformation, using the Iterative Closest Point algorithm (ICP) (Besl & McKay, 1992), and the residual distance d_A→B_ between the two sulci after registration was captured using the Wasserstein distance (Dobrushin, 1970). This choice of distance differed from the original approach (Sun et al., 2016) where the residual distance was captured using directly the distance from the ICP algorithm. In this study, this was a problem due to the fact that we were manipulating sulci of different size, in spite of the affine normalization to the Talairach space. Indeed, the length of the central sulcus relative to the brain size differs between the two time steps. Hence, using the ICP method, to register sulcus A on sulcus B, each point of sulcus A was matched to a point of sulcus B, but not every point of sulcus B was assured to have a match from sulcus A (as some points from sulcus B have been ignored for the registration). The residual distance minimized by the ICP registration was the sum of distances between the points in sulcus A and their match from sulcus B. This was satisfactory for the registration, but not to capture the dissimilarity between the two sulci, because it would have underestimated the distance when registering a small sulcus to a big one (e.g. if only the lower half of sulcus B exactly matched the point cloud from a shorter sulcus A, the upper half of sulcus B would have been completely ignored and the resulting distance would have been zero, even though sulcus A and B actually differed geometrically). The Wasserstein distance solved this problem by capturing the distance between the whole set of points from both sulci. Thus, we used the ICP algorithm for registration and then captured the resulting Wasserstein distance as the dissimilarity metric using the POT python toolbox (Flamary & Courty, 2017). After aligning sulcus A to sulcus B and capturing d_A→B_, sulcus B was registered to sulcus A using the same method, resulting in the residual distance d_B→A_. To mitigate the effect of a potential poor registration, the resulting dissimilarity metric was chosen as d _A,B_ = min(d_A→B_, d_B→A_).

### Use of distance matrix as shape descriptor for the whole cohort

This allowed us to build a 284x284 dissimilarity matrix, capturing the shape variability of the central sulci over the whole cohort through pairwise distances between sulci. In order to capture the main shape features of the whole cohort, we chose to operate a non-linear dimension reduction algorithm, the Isomap (Tenenbaum, 2000), which is detailed in the next section.

These first steps are summarized on Figure 2.

**Figure 2.**
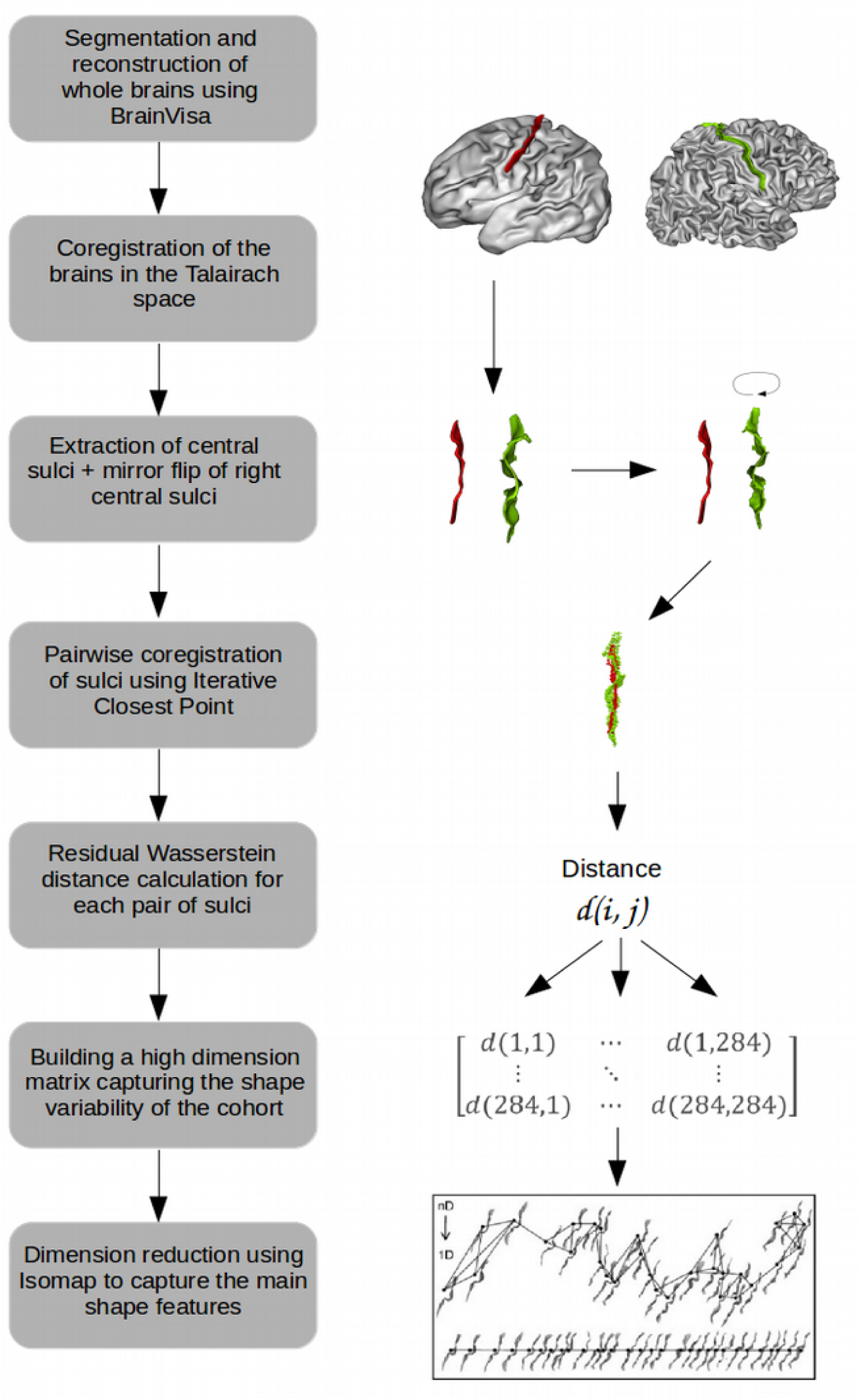
Summary of the preparatory steps for shape characterization.

### Choice of parameters for the Isomap algorithm

The Isomap algorithm builds a matrix of geodesic pairwise distances using an input dissimilarity matrix, by constructing a graph linking each sulcus to its *k* nearest neighbors, with *k* an adjustable parameter. The geodesic distance between any pair of sulci is then defined as the sum of the distances traveled by following the shortest path in the graph. A Multi-Dimensional Scaling algorithm is then applied to this geodesic distance matrix, to project the input data (the sulci) on a lower-dimension space of dimension *d*. Practically, the output consists of the coordinates of each sulcus on *d* axes representative of each dimension.

This algorithm requires two parameters: the number of nearest neighbors *k* used to build the neighborhood graph, and the number of dimensions *d* on which to project the geodesic distance matrix. The methods used to choose the *k* and *d* parameters are detailed in Annex 1, and resulted in the choice of *k* = 11 and *d* = 10.

### Visualization of the shape characteristics (descriptive approach)

To help interpreting the shape characteristics encoded by each selected Isomap dimension computed from the whole cohort, we used moving averages of the embedded sulci. In order to maximize the readability of these moving averages, we separated the sulci at 30w PMA from those at 40w PMA. This was necessary since the 30w PMA sulci are less developed than the 40w ones, and averaging both age-groups together for visualization would thus mix flatter and curvier sulci, making the shape interpretation complex. After separating the groups, the first step was to project the sulci on an axis, based on their Isomap coordinates for each dimension, after a rigid alignment of each sulcus to the most neutral one, namely the sulcus minimizing the average distance to the whole set. This resulted in *d* axes with the sulci ordered by shape characteristics. As this visualization was difficult to interpret, we used moving averages computed for a set of regularly spaced coordinates to help us identify the shapes encoded for each dimension. The range for the shape coordinates was defined so that at least 10% of the sulci at each time point (n=14) could be closest to the extreme coordinates, in order to prevent from building moving averages from very few sparse sulci on the borders of the dimensions. Each moving average was constructed by weighting the sulci (represented as point clouds) depending on their distance to the moving average coordinate, then summing them, then convolving the sum with a 3D Gaussian, and finally thresholding the result. The resulting shape was an average of the sulci’s shape in close vicinity to its location. We chose to generate ten equidistant medium shapes on each dimension. Once the moving averages were generated at both ages, we used the 40w PMA moving averages to interpret the shape characteristics encoded on each dimension. A reading key for shape description and a dimension description resulting from this method is illustrated on Figure 3.

**Figure 3.**
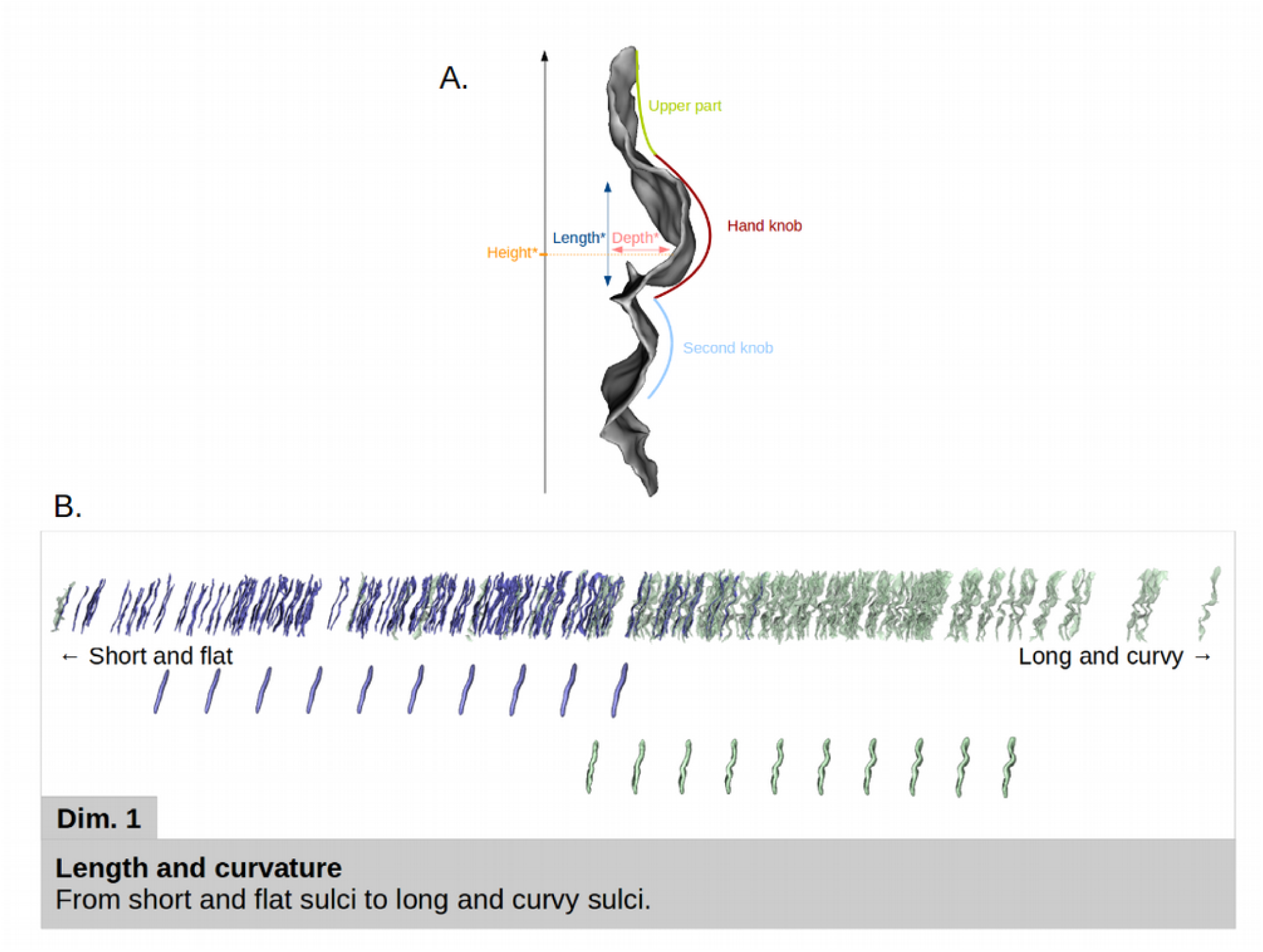
A: Reading key for shape description of the central sulcus with a two-knob configuration. The height of the hand-knob is defined as the vertical distance from the bottom of the sulcus to the deepest region of the hand-knob. B: Representation of the first Isomap dimension. In blue (resp. green): 30w PMA (resp. 40w PMA) sulci and moving averages. Top: sulcal projection. Middle: 30w PMA moving averages. Bottom: 40w PMA moving averages. Simple shape descriptors are given under the sulcal projection, more detailed descriptors are given in the lower box.

### Analyses of the developmental evolution (comparative approach of the two age-groups)

### Correction for age discrepancy at MRI acquisition

We hypothesized that the PMA variability around each MRI acquisition time-point could bias the shape characteristics. Therefore, we tested if the coordinates for any Isomap dimension correlated with PMA at acquisition, independently for each age-group but together for left and right sulci (see Sup. Info., Table 2). The resultant correlations (going up to 0.30 for dimension 1 at 30w PMA) confirmed the importance of correcting the dimension coordinates for PMA at MRI acquisition in each group, before conducting further analyses.

**Table 2.** Cross-validated scores for fine motor outcome classification. Scores obtained using 1) clinical factors; 2) sulcal shape features (positions corrected for PMA on the 10 dimensions), for each age and side subgroup; and 3) combination of both (clinical factors and sulcal shape features), for the fine motor outcome discrimination at 5 years of age. The bold values are the ones equal or superior to the baseline, i. e. the values obtained for the clinical factors alone.

|  | Subgroup | Balanced accuracy | ROC AUC |
| --- | --- | --- | --- |
| 1) Clinical factors (baseline) |  | 0.58 | 0.59 |
| 2) Sulcal shape alone | L30 | **0.62** | **0.66** |
|  | R30 | 0.45 | 0.44 |
|  | L40 | 0.40 | 0.38 |
|  | R40 | **0.62** | **0.66** |
| 3) Combination of clinical factors and sulcal shape | L30 | **0.59** | **0.61** |
|  | R30 | 0.48 | 0.50 |
|  | L40 | 0.46 | 0.41 |
|  | R40 | 0.57 | **0.64** |

ROC AUC: Area Under the Receiving Operator Curve. L: left hemisphere; R: right hemisphere. 30: 30w PMA; 40: 40w PMA.

For this purpose, we decided to perform the quantitative analyses on the Isomap dimensions after correction for PMA in each group: instead of using the raw Isomap coordinates for each dimension as the input for the subsequent analyses, we used their residuals after correction for PMA (that we centered on the age-group mean position on the Isomap in order to restore the information about inter-age-group positioning). The residuals were computed by solving, independently for each age-group, the linear model:

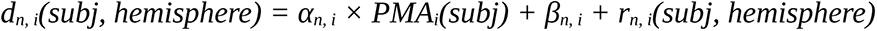

with *d_n, i_* the raw position of the sulci from the i-week acquisition (i= 30 or 40) on the n^th^ dimension of the Isomap, PMA_i_ the post-menstrual age at the i-week acquisition, *α_n, i_* and *β_n, i_* constants estimated by the model, and *rn, i* the sulci’s residual position on the n^th^ dimension after correction for PMA at the i-week acquisition.

Therefore, the shape characteristics considered in the following analyses, and which we refer to as Isomap positions corrected for PMA, were defined as:

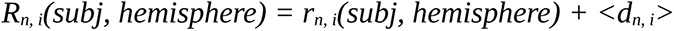

with <dn,i> the mean position of the sulci from the I-week acquisition (i= 30 or 40) on the n^th^ dimension of the Isomap.

### Comparison of the subjects’ positioning on the different dimensions

In the following, individual sulci from the two age-groups (i.e. 30w and 40w acquisitions) were considered separately, as well as left (L) and right (R) sulci. For each Isomap dimension, we investigated the relative positioning of the subgroups using two tests: a Wilcoxon signed-rank test to investigate whether the two age- or hemisphere-groups differed in positioning on a specific dimension (suggesting a difference in the shape of the sulci throughout development or between hemispheres), and a Spearman correlation to assess whether the two age- or hemisphere-groups showed correlated shape features (suggesting early and stable shape patterns throughout development or between the hemispheres). We conducted these tests between ages for each hemisphere (L30 vs L40, R30 vs R40), and between hemispheres at each age (L30 vs R30, L40 vs R40). Applying a correction for multiple comparisons with the Bonferroni approach would have led to a statistical threshold at 0.00125 for an alpha level at 0.05 because of the 40 tests performed (4 age-and-hemisphere-specific tests x 10 dimensions tested). However, as the Bonferroni method may be considered as too restrictive, we focused on results with p-values under the threshold computed for each age-and-hemisphere-specific test, thus only compensating for the number of dimensions, resulting in a statistical threshold at 0.005. Only the results with p-values below or in the same range as this corrected threshold are described in the Results section. The whole test results are available in Sup. Info., Table 3 for Wilcoxon signed rank tests and Table 4 for Spearman correlations.

### Handedness and fine motor outcome predictability

### Purpose of prediction of handedness and fine motor outcome

We aimed to predict the children’s handedness and fine motor outcome (manual dexterity score from the mABC scale) assessed at 5 years of age, based on the early shape of the central sulcus. We therefore performed two different analyses, aiming to classify either left versus right handers, or children with poor versus good fine motor outcome, using three sets of features: 1) a selection of early clinical factors (without sulcal considerations), 2) the shape features of the central sulci (without clinical considerations), encoded by the Isomap positions corrected for PMA, and 3) the combination of both. The purpose of using these three sets of features was to be able to compare the predictive capacity of these different features, and more specifically to assess whether sulcal shape is a better predictor for motor development than early clinical factors, and whether the combination of both outperforms the two previous classifiers, suggesting a complementary influence of early morphological and clinical features on the long-term motor development.

### Dichotomization of the motor classes

The cohort was dichotomized for each motor characteristic: right versus left handedness, and poor versus good fine motor outcome (manual dexterity).

Children born preterm display a significantly higher occurrence of non-right handedness than full-term children (odds ratio 2.12) according to a meta-analysis (Domellöf et al., 2011). Studying handedness in preterm children may be a potentially important index reflecting hemispheric organization and sensorimotor functions ensuing neurodevelopmental disturbances. Thus, we hypothesised that among the left-handed children in our cohort, some would be “natural” left-handers while the others may express a non-right-hand preference following a disruption in the lateralized early brain organization associated with premature birth. Since a previous study reported links between handedness and the shape of the central sulcus on healthy adults (Sun et al., 2012), we decided to focus on typical handedness by maximizing the proportion of “natural” left-handers (n=7 left-handers with at least one left-handed parent (Bryden et al., 1997), subsequently referred to as “left-handers”) to compare with right-handed (n=50) children. We thus excluded children with missing data (n=1), ambidextrous children (n=2) and left-handers for whom both parents were right-handers (n=11).

In terms of fine motor assessment provided by the mABC score, we focused on the manual dexterity subscore, which assessed three tasks: posting coins, threading beads, and drawing a trail. We chose to focus on this score because links between functional activation of the hand and the shape of the central sulcus (more specifically the height of the hand-knob) have already been reported in adults (Sun et al., 2016), and we therefore suspected that, compared to the other two mABC subtests – aiming/catching and balance – it was the subtest with the most direct link to the central sulcus’s shape.

We dichotomized the children group according to their poor and good fine motor outcome, removing children with borderline results from the classification. For the mABC total score, a score inferior or equal to the 5^th^ percentile indicates definite motor problems; between the 6^th^ and 15^th^ percentiles indicates borderline performance; and strictly above the 15^th^ percentile indicates normal motor development (van Heerwaarde et al., 2020). Since for the manual dexterity subscore no percentile cut-offs have been assessed, we used the same cut-offs in our study population, with the corresponding standard scores for each percentile range: children whose standard score ranked between 1 and 5 were assigned to the poor fine motor outcome group (n=15), children who ranked strictly above 7 were assigned to the good fine motor outcome group (n=35), and children ranking 6 or 7 (n=16) were considered as borderline and were not included in the classification analyses.

### Subgroup differentiation and feature choice

In terms of clinical factors, the retained variables were the following: gestational age at birth, the birth-weight z-score, the presence of a severe intra-ventricular hemorrhage, and the presence of broncho-pulmonar dysplasia. All the continuous variables were normalized, and the categorical variables were binary, encoded as 0 or 1.

In terms of shape features of the central sulcus, the features retained were the Isomap positions corrected for PMA at scan, on the 10 relevant Isomap dimensions. We once again considered separately the L30, R30, L40 and R40 subgroups, because of the high variability in sulcal shapes at both ages, and because we expected that inter-hemispheric asymmetries in shape might play a role on handedness and fine motor outcome.

The classifiers were trained once per age and side subgroup, except for the classifiers using the clinical factors as only features, since the clinical factors considered are neither age nor side dependent.

### Cross-validated training of subgroup-specific classifiers

We used a linear Support Vector Classifier (SVC) to assess the predictive capacity of the shape and clinical features on the handedness and fine motor outcome. We parametrized the SVC to take into account the imbalance between classes (either 7 vs 50 or 15 vs 35) by weighting the regularization parameter by the inverse of class frequency in the input data. Due to the small size of the groups (either 57 or 50 sulci for handedness or fine motor outcome respectively), we took two precautions to limit the chances of overfitting our model: we used the predetermined regularization value (C=1), and we used a Stratified Shuffled Cross-Validation (number of folds: 5, number of repeats: 10) to evaluate the performance of the algorithm. We then evaluated the balanced accuracy (the average of correctly predicted samples for both classes) and Area Under the Receiving Operator Curve (ROC AUC). For the best classifier using only sulcal shape for prediction, we plotted the ROC and Precision-Recall (PR) curves, next to the ROC and PR curves obtained for the clinical factors alone for comparison.

### Observation of most-weighted features

For the classifiers using only sulcal shape which showed the best performance, we were interested in the most relevant shape features in the training of the classifiers. Therefore, during the cross-validation, we stored the coefficients attributed to each Isomap dimension. After the cross-validation, we looked into the consistency of coefficient attribution depending on the training/testing set using a box plot picturing the coefficient attributed to each Isomap dimension for every iteration of the cross-validation. More importantly, we assessed which shape features were the most discriminative for each outcome. The dimensions corresponding to these specific shape features were described in more detail.

## Results

### Characterization of the central sulcus’ shape Visual Isomap results

For brevity concerns, we only present, in the following sections, the visual representations of the dimensions relevant in the performed analyses. The detailed descriptions for all 10 dimensions (not corrected for age, to improve interpretation) are shown in Annex 2.

### Age dependency of Isomap positioning

The Pearson correlation coefficients between age at MRI acquisition and raw Isomap position (not corrected for age) are reported for the 30w PMA and 40w PMA sulci separately in Sup. Info., Table 2. The highest correlation observed was r=0.30 for the first Isomap dimension at 30w PMA, the dimension coding for length and global curvature of the sulci. Along with the other coefficients observed, the correlations suggested that the Isomap position of sulci is somewhat affected by PMA, which justified our decision to correct for it in subsequent analyses.

### Hemispheric comparisons for each age-group

The Wilcoxon signed-rank test results obtained by comparing the PMA-corrected positioning of left and right sulci on each Isomap dimension, for each 30w and 40w PMA group, highlighted in Sup. Info., Table 3.A. two dimensions (one per age-group) with significant hemispheric asymmetry (Sup. Info., Table 3.A, Figure 4). At 30w PMA, dimension 3 (coding for the curvature of the hand-knob at fixed height) suggested that left central sulci were roughly flatter than the right ones. At 40w PMA, dimension 8 (coding for the switch from a single to a double-knob configuration) also showed some asymmetry, with more left central sulci in the double knob configuration and more right central sulci in the single knob configuration.

**Figure 4.**
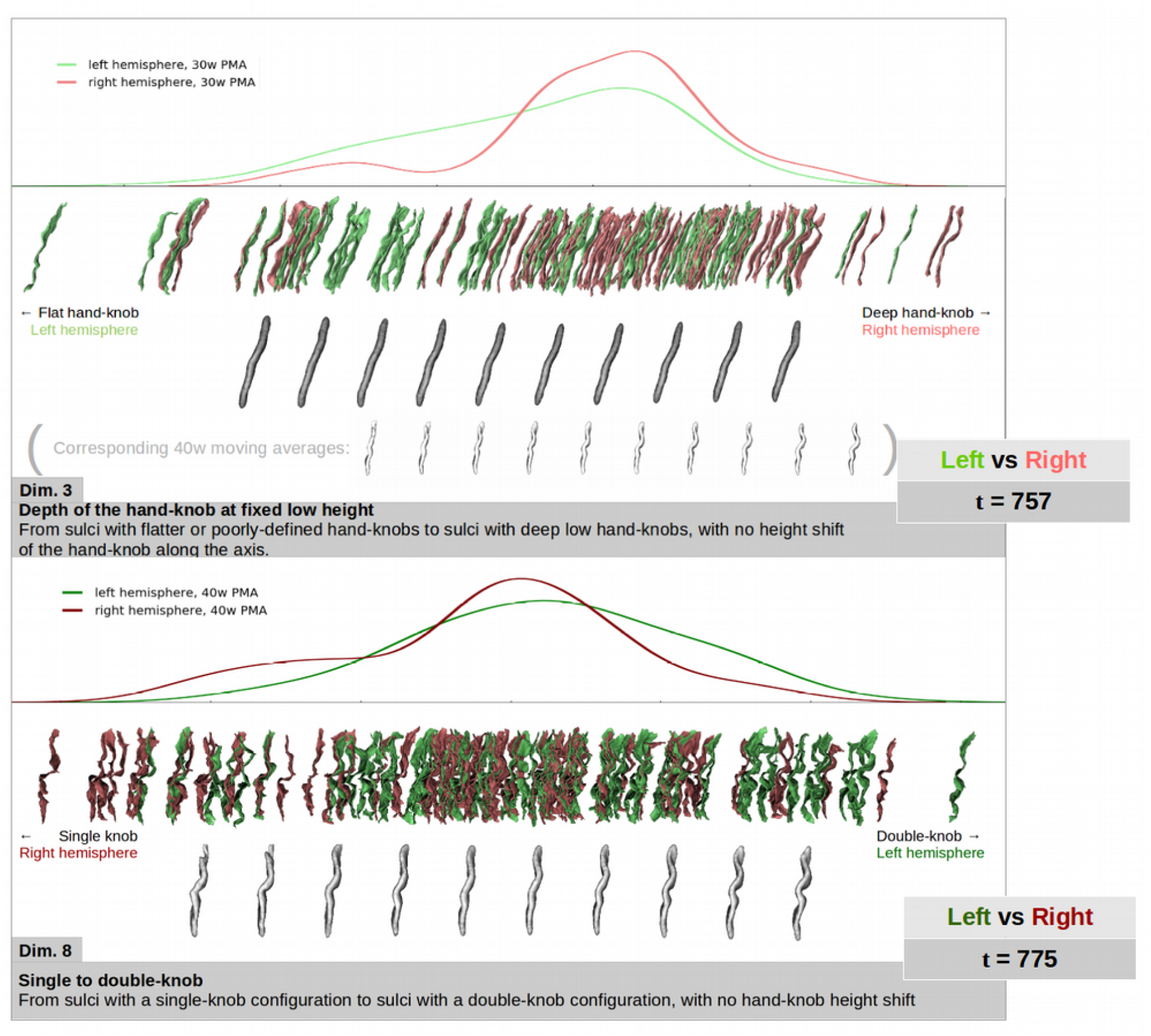
Representation of left (green) versus right (red) central sulci distribution on the two dimensions showing statistically significant hemispheric asymmetries (Dim. 3 and 8). Upper rows: density plot of sulcal distribution along the dimension. Lower rows: sulcal projection and corresponding moving averages (in grey). On the 30w PMA representation of Dim. 3, the 40w moving averages (in white) are shown for shape interpretation purposes.

In terms of Spearman correlations (Sup. Info., Table 4.A), four dimensions showed a significant correlation between the left and right central sulci: dimension 1 at both ages, dimension 5 and 2 at 30w PMA, and dimension 3 at 40w PMA. These results suggest that at 30w PMA, left and right central sulci show a coherent encoding of length, curvature (dimension 1), wrapping around (dimension 5) and height (dimension 2) of the hand-knob, and depth of the second knob (dimension 2). At 40w PMA, left and right central sulci seem to show consistent sulcal length (dimension 1) and depth of the hand-knob at low fixed height (dimension 3).

### Age-group comparisons of the hemisphere-specific shape characteristics

Wilcoxon signed-rank tests performed to compare PMA central sulci characteristics (Sup. Info., Table 3.B) identified several dimensions with different positions at 30 versus 40w (Sup. Info., Table 3.B). Two dimensions (1 and 5) were highlighted on both hemispheres, suggesting that sulci on both hemispheres are roughly shorter, flatter, and show a more wrapped-around hand-knob at 30w PMA compared to 40w PMA. Two other dimensions were relevant only for one hemisphere: dimension 4 for the left hemisphere, with shallower and lower hand-knob at 40w than at 30w, and dimension 2 for the right hemisphere, with a higher hand-knob and a deeper second knob at 40w PMA compared to 30w PMA.

Out of the 10 dimensions of interest, we observed relevant trends in Spearman correlation between the PMA-corrected Isomap positioning at 30w and 40w PMA on 8 dimensions (Sup. Info., Table 4.B). The three dimensions showing the strongest correlations between both ages are presented in Figure 5. Dimension 4 was observed on both hemispheres and encoded a variation of height and depth of the hand-knob, as well as the presence and depth of the second knob (i.e. from double-knobbed sulci with deeper and higher hand-knobs, to sulci with a single, lower, and shallower hand-knob). On the left hemisphere, dimension 2 showed correlations between 30w and 40w PMA in the height of the hand-knob and the depth of the second knob. On the right hemisphere, the highest correlation was observed on dimension 8, correlating the sulci depending on their single-knob or double-knob configurations, with no height shift of the hand-knob. This suggested that between 30 and 40w PMA, most of the early morphological characteristics (8 dimensions over 10) tend to evolve into a more complex but consistent encoding (i.e. a sulcus with a single, low and shallow hand-knob at 30w will most often show the same characteristics at 40w, rather than a double-knob configuration).

**Figure 5.**
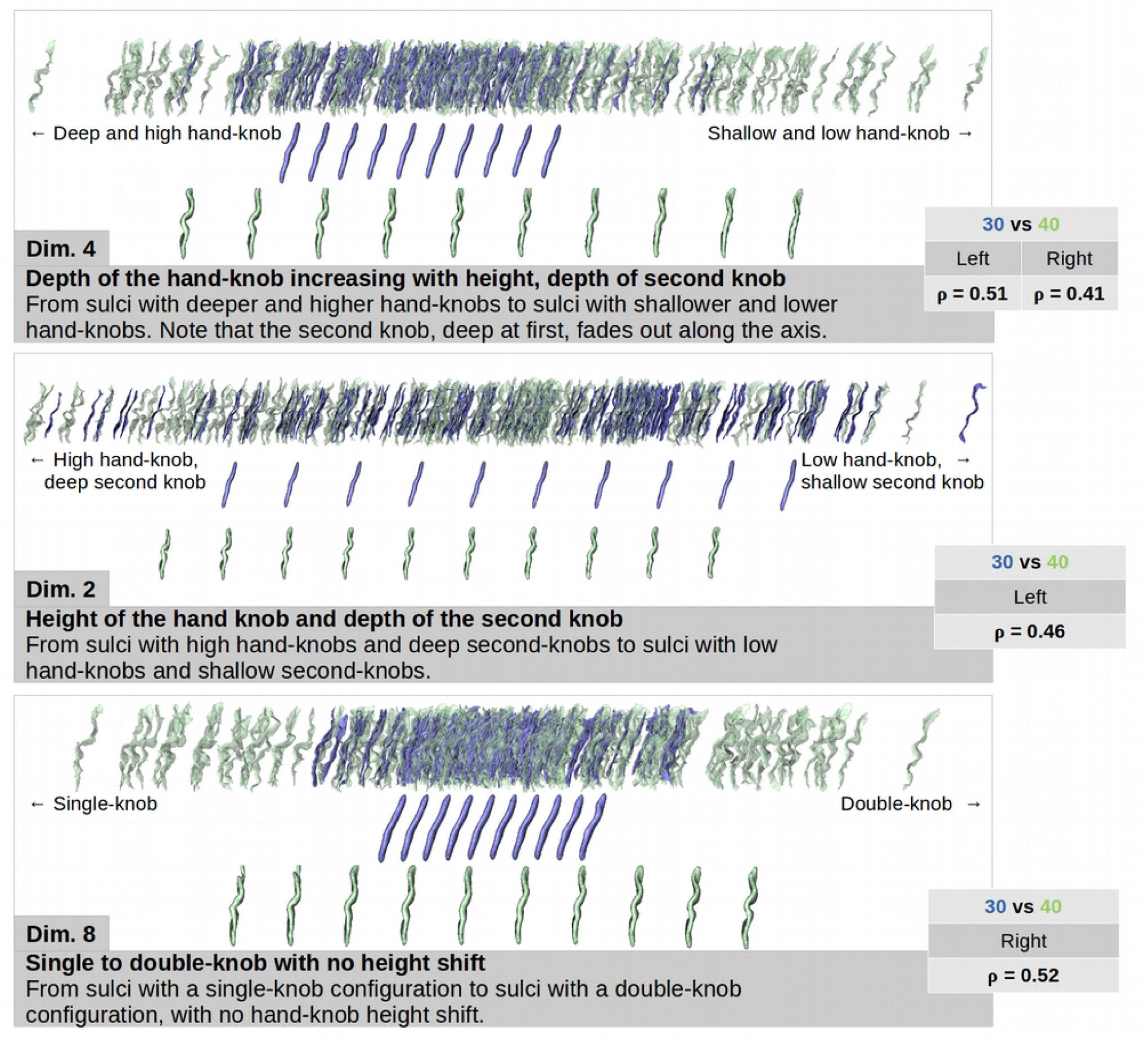
Representation of the three dimensions showing the strongest correlations between 30w (blue) and 40w (green) PMA central sulci. For each dimension, the sulcal projections were represented in the upper rows, and the age-specific moving averages on the lower rows .

This suggested that between 30 and 40w PMA, most of the early morphological characteristics (8 dimensions over 10) tend to evolve into a more complex but consistent encoding (i.e. a sulcus with a single, low and shallow hand-knob at 30w will most often show the same characteristics at 40w, rather than a double-knob configuration).

### Classification of handedness and fine motor scores based on sulcal shape features

### Discrimination by shape for left- vs right-handers: classifiers’ performance

The support vector classifiers were trained to differentiate right-handers (n=50) from left-handers (n=7). The cross-validated scores of the classifiers considering either clinical factors, age- or hemisphere-specific shape features (corrected for PMA) or both, are shown in Table 1.

The best classifier (highest balanced accuracy and ROC AUC) was obtained using the combination of clinical factors and sulcal shape features at 30w PMA on the right hemisphere. Yet, the performances obtained with these shape features alone were quite low. In contrast, the best classifier using only shape features was the one obtained at 30w PMA on the left hemisphere, and it exceeded the performance of the classifier using only clinical factors, while adding the clinical factors to these shape features also provided a quite relevant classifier. That is why we further focused on the classifier obtained with shape features of left sulci at 30w PMA alone. The ROC and PR curves of this classifier (and correspondent curves obtained for clinical factors alone as a comparison) are shown in Figure 6.

**Figure 6.**
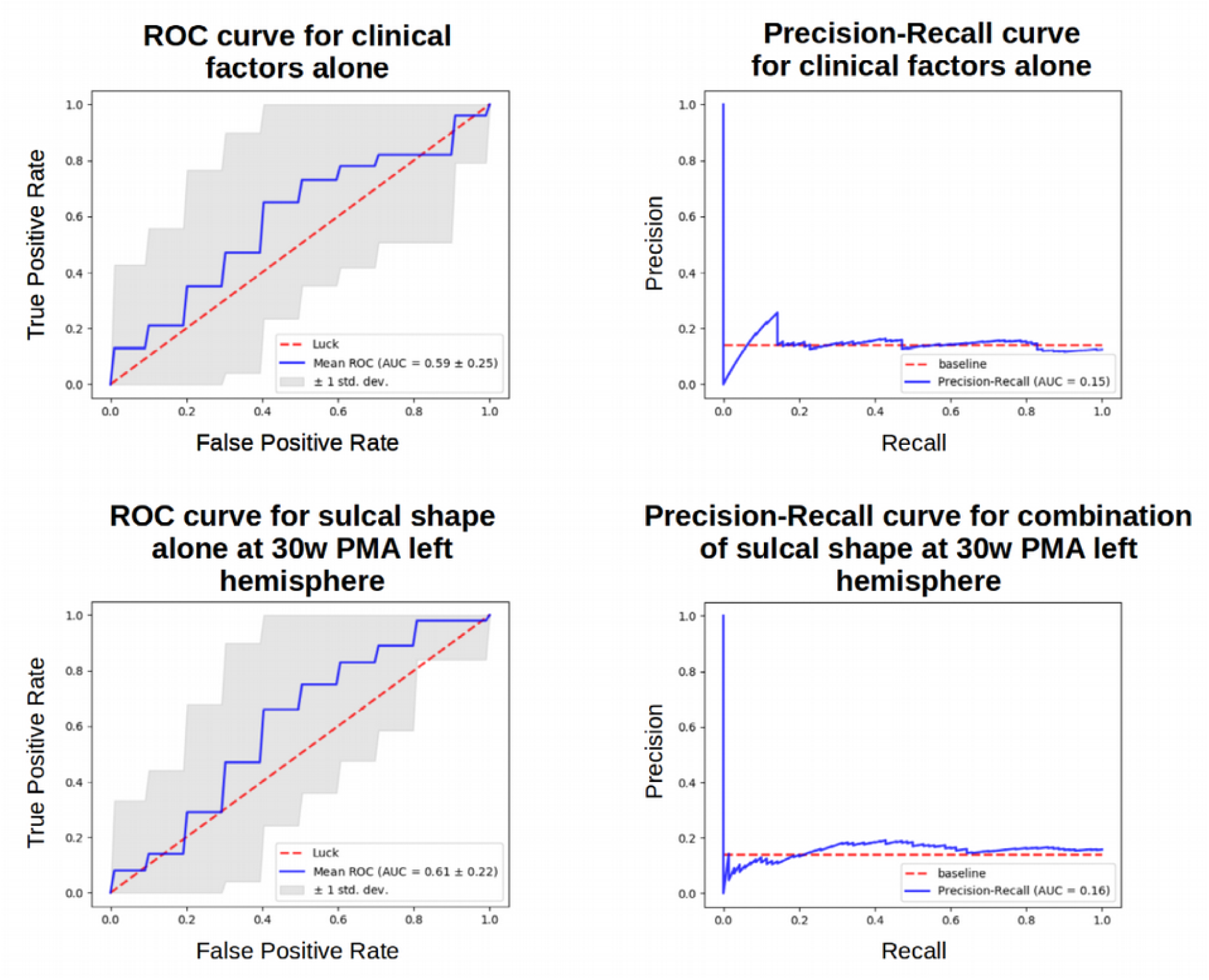
ROC and PR curves obtained for clinical factors alone and sulcal shape factors alone at 30w PMA on the left hemisphere. Target: handedness

For this classifier, we further wondered which shape features were considered relevant in differentiating left-handers and right-handers. Using the different folds of the repeated stratified cross-correlation, we aggregated the coefficients attributed to each Isomap dimension and generated the box plot presented in Figure 7.

**Figure 7.**
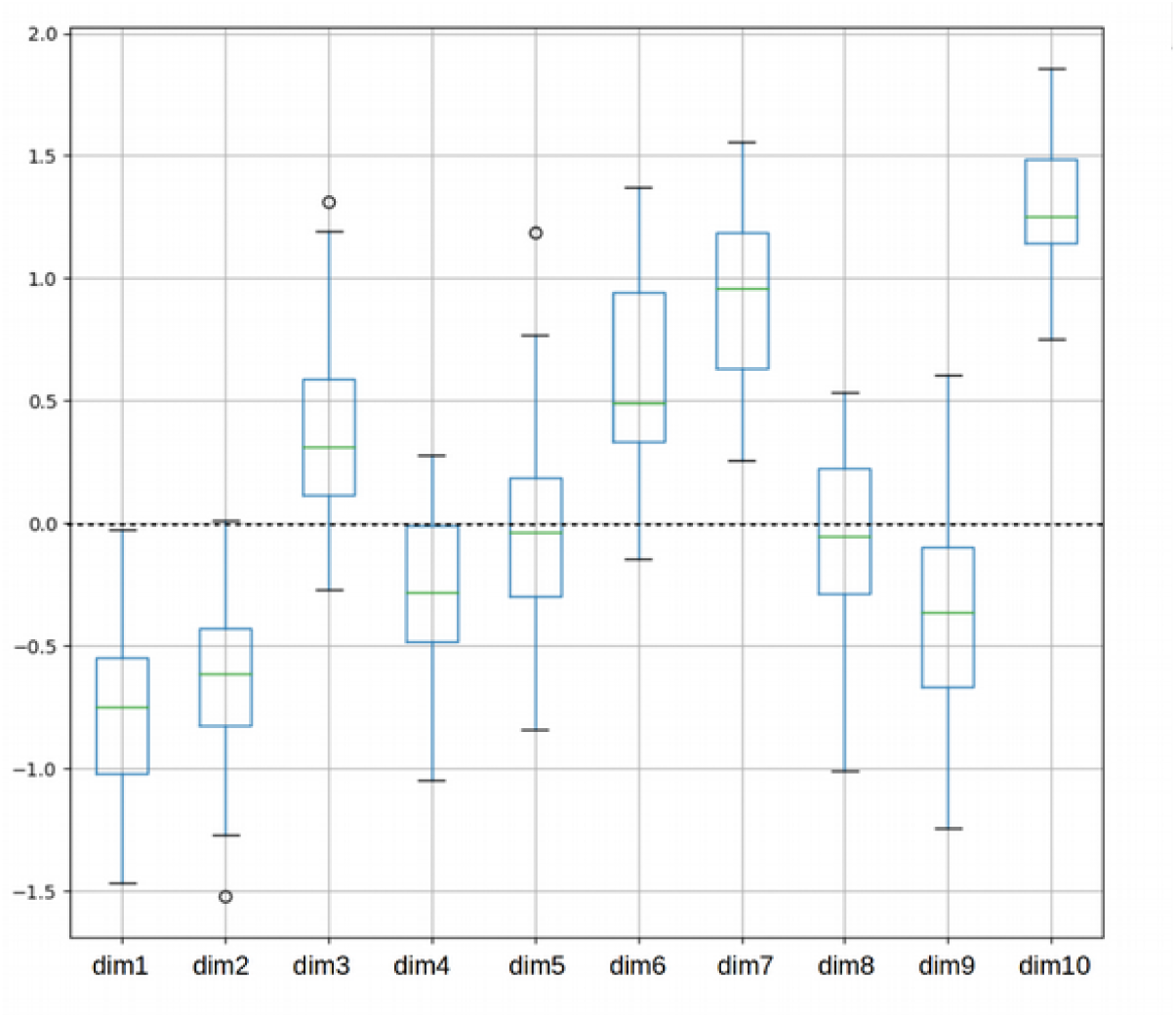
Box plot of the coefficients attributed to each Isomap dimension during the cross validation for the handedness classification with sulcal shape factors alone at 30w PMA on the left hemisphere. Coefficients for the dimensions were obtained for each iteration of a 5-fold 10-times repeated stratified cross-validation training of the SVC for handedness. Dimension 10 weighted generally more than the other dimensions throughout this cross-validation.

Most Isomap dimensions were reasonably consistent in weight through the cross-validation. The 10^th^ dimension was clearly and almost systematically the most informative. The visual interpretation of this dimension (Figure 8) suggested that the most discriminative traits for handedness were the length of the hand-knob and the orientation of its upper part. Compared to right-handers, the left-handers tended to have a left central sulcus at 30w PMA with a longer hand-knob, with its upper part bending backwards.

**Figure 8.**
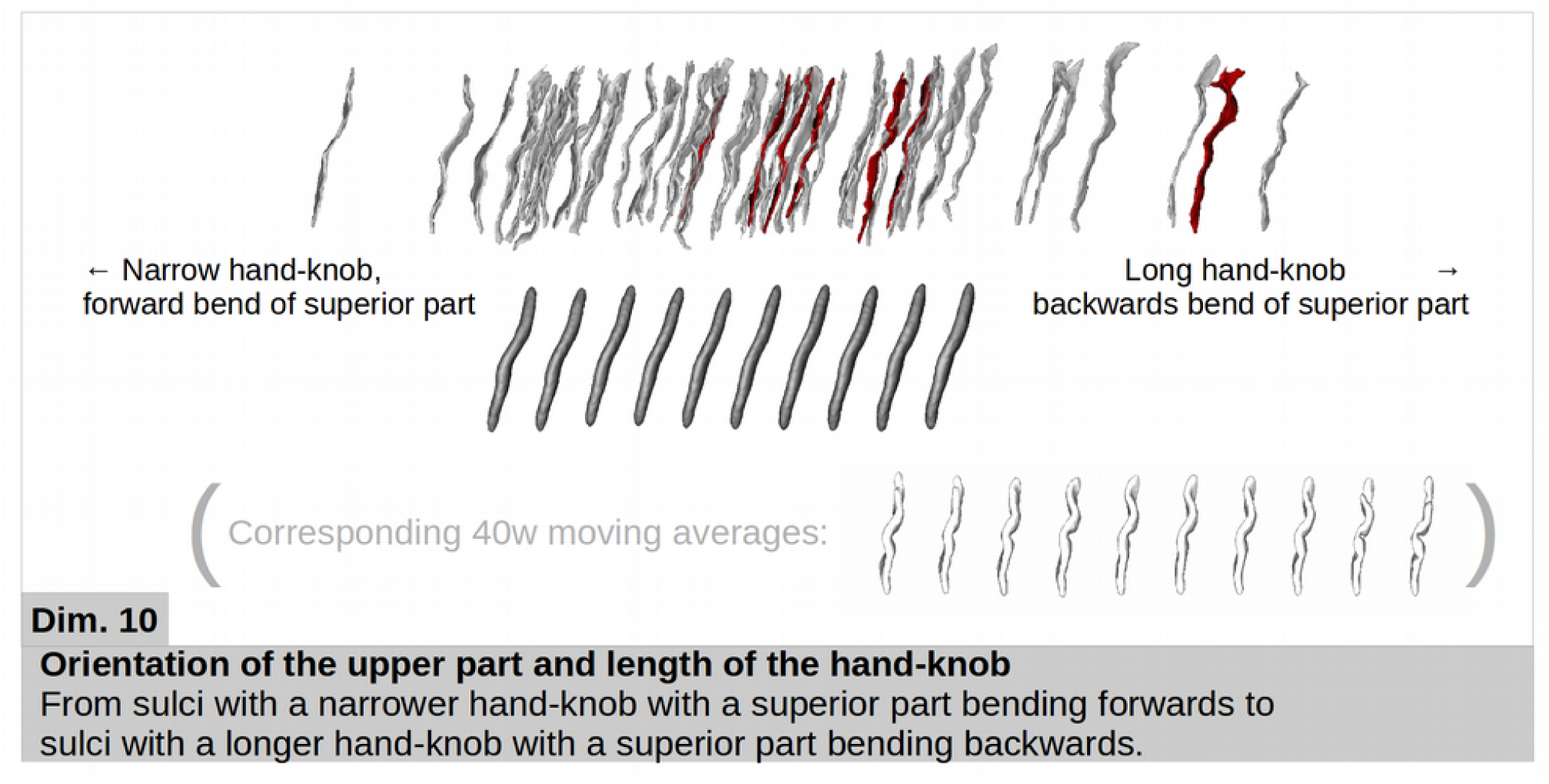
Representation of the dimension weighting the most for handedness classification using sulcal features at 30w PMA in the left hemisphere. The sulcal projections of the subjects used in the classification were represented (right handers in grey and left handers in red), as well as the age-specific moving averages (in dark grey). The 40w PMA moving averages (in white) are also shown for shape interpretation purposes.

### Discrimination by shape for the fine motor outcome: classifiers’ performances

The support vector classifiers were trained to differentiate subjects with poor fine motor outcome (n=15) from subjects with good motor outcome (n=35), again considering either clinical factors, age-and-hemisphere specific shape features (corrected for PMA) or both. The cross-validated scores of the classifiers are shown in Table 2.

Two classifiers scored a tie for balanced accuracy and ROC AUC, using the sulcal shape features alone: the left hemisphere at 30w PMA and the right hemisphere at 40w PMA. We chose to focus on the one with the best recall (proportion of correctly identified poor fine motor outcome, the most interesting clinical issue), which was the one for the right hemisphere at 40w PMA (recall = 0.61 against recall = 0.53 for 30w PMA left hemisphere). For this classifier, the ROC and PR curves are shown in Figure 9. The results for the classifier obtained at 30w PMA on the left hemisphere using only the sulcal shape features are presented in Annex 3.

**Figure 9.**
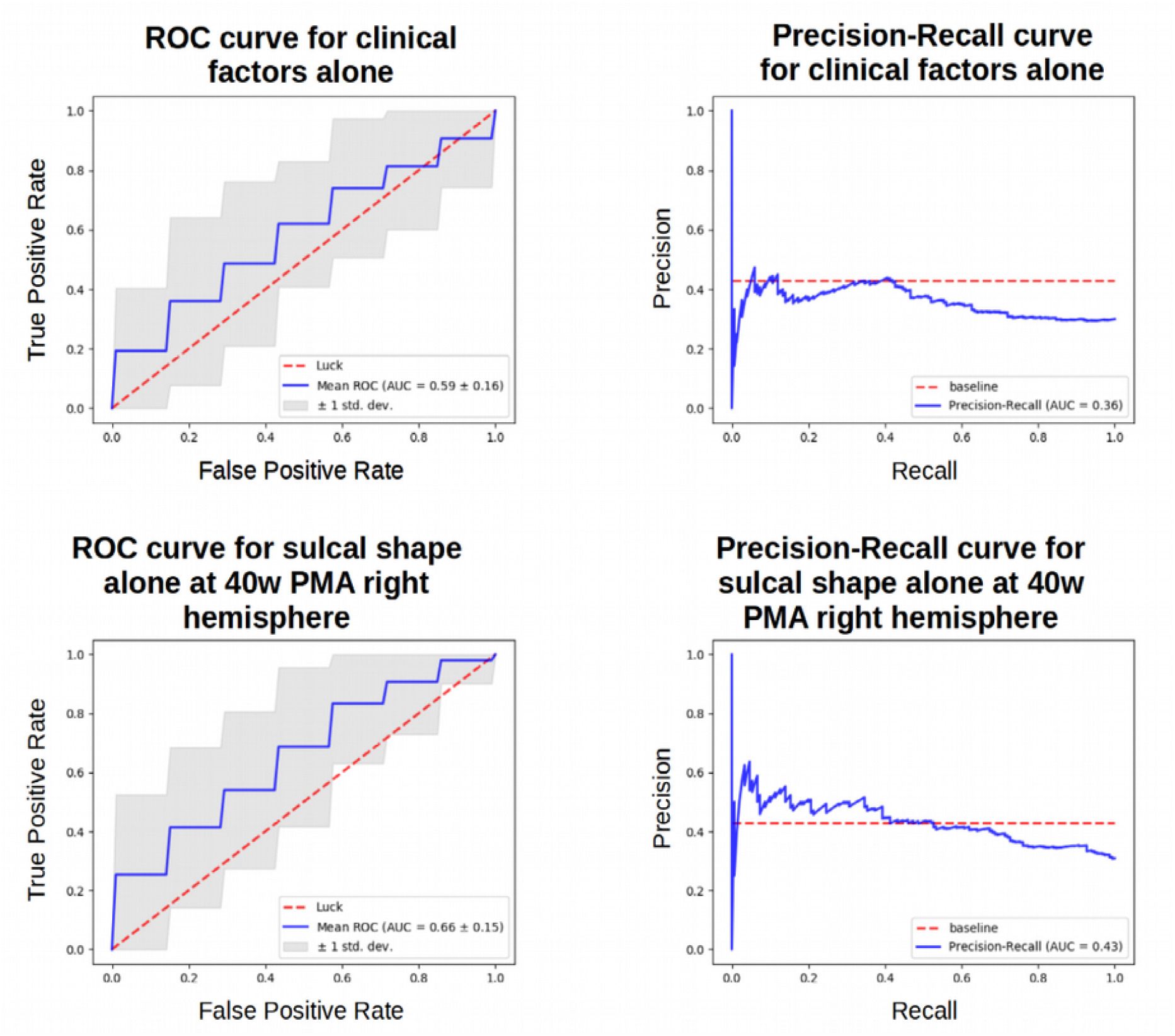
ROC and PR curves obtained for clinical factors alone and sulcal shape factors alone at 40w PMA on the right hemisphere. Target: fine motor outcome.

This latter resulting classifier performed better than a random classifier, and again we addressed the question of the relevance of the different shape features in the classification. Using the different folds of the repeated stratified cross-correlation, we aggregated the coefficients attributed to each feature and generated the box plot presented in Figure 10.

**Figure 10.**
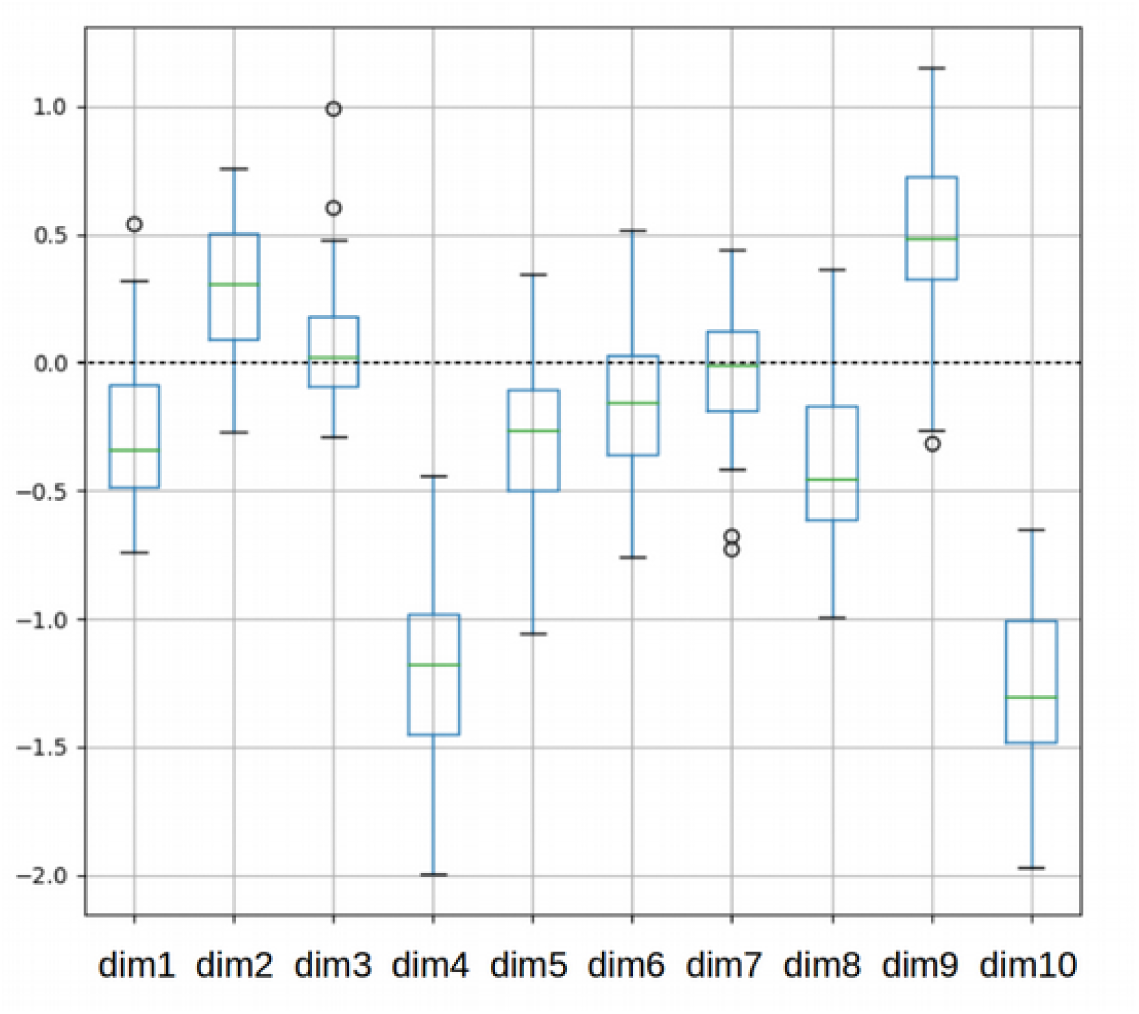
Box plot of the coefficients attributed to each Isomap dimension during the cross validation for the fine motor outcome classification with sulcal shape factors alone at 40w PMA on the right hemisphere. Coefficients for the dimensions were obtained for each iteration of a 5-fold 10-times repeated stratified cross-validation training of the SVC for fine motor outcome. Dimensions 4 and 10 weighted generally more than the other dimensions throughout this cross-validation.

Once again, the dimensions were rather consistent in weight through the cross-validation. Dimension 10 was the most relevant dimension, but dimension 4 seemed to have a comparable role in the classification, so we considered both dimensions for the shape description (Figure 11). The visual interpretation of these dimensions suggested that children presenting a poor fine motor outcome at 5 years of age tend to have a right central sulci at 40w PMA with a shorter but higher and deeper hand-knob, with a superior sulcal part leaning frontally.

**Figure 11.**
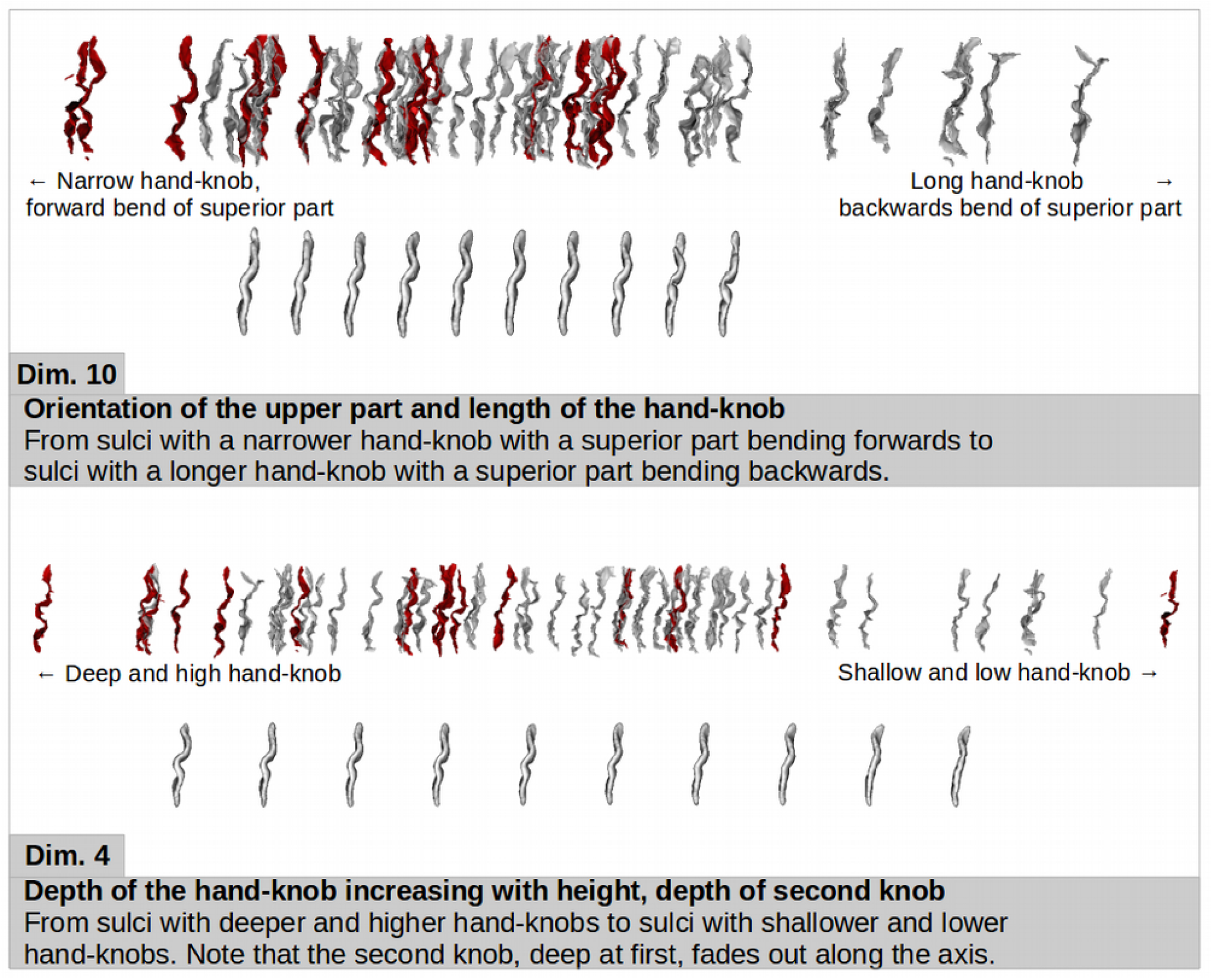
Representation of the two dimensions weighting the most for fine motor outcome in the classification at 40w PMA in the right hemisphere. For each dimension, the sulcal projections of the subjects used in the classification were represented (good outcome in grey, poor outcome in red), as well as the age-specific moving averages (lower row, in light grey).

### Discussion

This study 1) longitudinally described the shape characteristics and variability of the central sulcus in extremely preterm infants at 30 and 40w PMA; 2) brought to light hemispheric asymmetries at each age; 3) showed a close developmental relationship between the sulcal shape at both postnatal ages; and 4) unraveled links between the perinatal shape of the central sulcus and the later handedness and fine motor outcome in this cohort of children born extremely preterm.

### How to study cortical folding

Different approaches have been developed and used to study the *complexity and implications of cortical folding*, but few have described its variability, and even fewer specifically in the developing brain. Sulcal information has been collected through various means and for multiple purposes. The simpler sulcal information includes basic geometric measures, such as length, surface area or mean depth after sulcal extraction, as proxies for shape (Kersbergen et al., 2016). More sophisticated ones include the use of mean depth profiles (Gajawelli et al., 2021), the calculation of local or global gyrification indices (Cachia et al., 2008) or focus on sulcal pits (Régis et al., 2005; Im & Grant, 2019). Such studies addressed general brain development issues, such as normal development in the fetus (Vasung et al., 2019) or in the preterm newborn (Moeskops et al., 2015; Orasanu et al., 2016; Garcia et al., 2018), and more specific adversarial development questions, such as early neurodevelopmental events linked to pathological outcomes (Mellerio et al., 2015). Compared with previous studies, the approach developed in this article additionally describes and quantifies the inter-individual sulcal variability.

In relation to this issue of inter-subject variability, the historical method relied on visual examination and description of sulcal morphology (Ono et al., 1990) but implied long, subjective and potentially biased examination processes, and resulted in small sample sizes. Newer methods have been developed to automate both sulcal extraction and shape characterization. They can either use a clustering approach or a continuous approach. Clustering methods try to identify a finite number of shape patterns to characterize a given sulcus. This approach seems appropriate for structures presenting great variability, such as the sulci from the anterior cingulate cortex (Del Maschio et al., 2019), or the gyri from different regions using multi-view curvature features (Duan et al., 2020). These clustering approaches are hardly suited for the study of the central sulcus because of the simplicity of its configuration: generally uninterrupted and without branches, except for a few individual exceptions (Mangin et al., 2019). Moreover, the clustering methodology is not suited for a longitudinal comparison of shape. Therefore, we opted for a continuous approach in our study. Some continuous approaches of sulcal shape description have relied on morphologic landmarks, such as the superior temporal asymmetrical pit in the superior temporal sulcus (Leroy et al., 2015), or the occipital bending of the Sylvian fissure (Hou et al., 2019). Both of these studies used these landmarks to quantify the difference in sulcation between humans and chimpanzees in order to highlight human specificities, and therefore did not focus on intra-species variability. Another continuous approach was the one previously used to study the central sulcal shape in adults (Sun et al., 2012; Sun et al., 2016). We chose to adapt this method to the developmental context of our study, in order to obtain a detailed characterization of the central sulcus’s shape, requiring neither prior assumptions about distinct clusters nor the definition of anatomical landmarks, as well as allowing multiple-age comparison.

The method retained, using pairwise coregistration of sulci to capture the shape dissimilarity within the cohort, allowed us to explore multiple shape characteristics of the central sulcus and therefore to capture its variability with unprecedented precision, leading to the observation of new specificities of this sulcus. In particular, the previous study assessing sulcal specificities on the same cohort using basic shape features reported a single significant hemispheric asymmetry in the central sulcus at 30w PMA – involving depth of the central sulcus but not surface area – and none at 40w PMA (Kersbergen et al., 2016). Our method, capturing multiple shape features, allowed us to reveal hemispheric asymmetries both at 30 and 40w PMA.

### How does preterm birth affect cortical folding around the central sulcus?

In order to capture the early folding dynamics of the developing brain, it would be ideal to study longitudinal data from fetal period to full-term birth, but using a mix of fetal and post-natal data is slippery because birth *per se* seems to affect the shape and folding of the brain (Lefèvre et al., 2016). Therefore, we chose to longitudinally study sulcal features in extremely preterm infants in order to investigate early brain development. Yet, prematurity has been reported to provoke alterations in sulcation: compared to term-born neonates, very preterm infants at term-equivalent age showed decreased sulcation index and folding power specifically in primary sulci (Shimony et al., 2016; Dubois et al., 2019), confirming that a very preterm population may not reflect perfectly typical folding of the central sulcus.

In this context, different studies have focused on the sulcation variability in the neonate. In particular, recent studies described the major sulcal or gyral patterns of cortical folding in newborns – including the central sulcus – using clustering approaches based either on deep sulcal pits (Meng et al., 2018) or on multi-view curvature features (Duan et al., 2020). Yet, to the best of our knowledge, the present study is the first to describe qualitatively and quantitatively the shape variability of the central sulcus in the preterm neonate, and more importantly to address the question of longitudinal shape evolution between early folding and the resulting sulcation pattern at term-equivalent-age.

In terms of qualitative morphological description of the central sulci, we presented a visual representation of the shape variability through 10 dimensions, and we concurrently suggested corresponding shape descriptions (see Annex 2). The scope of this work has yet to be explored in future studies: to what extent does the shape variability reported in this study describe globally the neonate population (including fetal and term-born populations)? What fraction of this variability is specifically associated either to prematurity or to related early pathological events? Since this first approach was exploratory, we decided to analyze the whole cohort regardless of clinical characteristics which can impact brain development (e.g. low gestational age, being a twin) (Dubois et al., 2008; Kersbergen et al., 2016) or other factors which might have a slight influence on our sulcal asymmetry observations (e.g. the positioning of the infant in the scanner: on the side at 30w PMA versus on the back at 40w PMA). To investigate the question further, it would be interesting to compare the difference in shape of the central sulcus between a preterm population at term-equivalent-age and healthy term-born subjects, in order to differentiate prematurity-related variability from typical variability. It would also be informative to investigate the effects of clinical conditions on early sulcation by differentiating subgroups of patients. Nevertheless, our cohort was too small to provide reliable results on this question. The application of our approach to other large databases, such as the developing Human Connectome Project (dHCP) (Makropoulos et al. 2018), is therefore an interesting prospect, but was beyond the scope of the current study.

In terms of quantitative shape description, we reported an early encoding of most shape features, with eight out of ten dimensions showing a statistically relevant correlation between 30 and 40w PMA. The fact that the early shape of the central sulcus remained mostly consistent despite the later development of secondary sulci – which mostly happens between 30w and 40w PMA (Dubois et al., 2019) – was informative about the interactions between folding waves. The secondary folding wave does not seem to drastically affect the central sulcus’s characteristics. To illustrate this, we could have suspected that the wrapping around the hand-knob (illustrated by dimension 5) might be affected by the secondary folding of the superior frontal sulcus (which might constraint the gyrus adjacent to the superior part of the central sulcus to retract dorsally and “push back” the upper part of the central sulcus, “unwrapping” the hand-knob). Yet, the correspondent Spearman correlation coefficients (0.38 and 0.32 for left and right hemisphere respectively) indicated that this feature evolved consistently between its 30w and 40w PMA encoding. The only two features showing low (ρ = 0.19) or very low (|ρ| < 0.1) correlations between the two age-groups on both hemispheres were dimensions 7 and 10, namely the height of the hand-knob at fixed depth and the orientation of the upper part and length of the hand-knob. These two specific features may have a delayed developmental dynamic or may present significant morphological changes between the two ages, which would explain why the 30w PMA encoding of these features did not foreshadow their 40w PMA counterparts. The knob height and the upper part orientation may depend on a shorter or longer period of elongation of the top of the central sulcus, which occurs between 30w and 40w PMA (Mangin et al., 2019).

Looking into the correlations between hemispheres, we also investigated hemispheric asymmetries in the central sulcus. A previous study on the same cohort reported solely a rightwards sulcal depth asymmetry at 30w PMA, and no hemispheric asymmetries in terms of surface area in the central sulcus, at both ages (Kersbergen et al., 2016). In this study, using more complex shape characterization techniques, we identified one dimension per age-group showing a statistically significant asymmetry. At 30w PMA, right central sulci showed a generally deeper hand-knob than left ones (dimension 3). This could be explained by the earlier maturation of the right hemisphere compared to the left one (Chiron et al., 1997), and by the earlier sulci formation in the right hemisphere (Chi et al., 1977; Dubois et al., 2008; Habas et al., 2012). Yet, with an earlier maturation of the right central sulci, we could have expected a length asymmetry, which would have been encoded on dimension 1 – which did not show a significant asymmetry. An interpretation would be that at 30w PMA the left central sulci have caught up to the right ones in terms of length, and the relative maturational advance of the right central sulci is therefore expressed in terms of depth of the hand knob rather than length of the sulcus. At 40w PMA, the left central sulci favoured a “double-knob” configuration (versus “single-knob” configuration for the right ones) (dimension 8). This finding is consistent with a previous study on adults, which reported a visually similar main shape variability (from single to double-knob configuration with hand-knob height increase), and asymmetry (Sun et al., 2012).

### To what extent is the central sulcal shape relevant to predict handedness and fine motor outcome?

A number of recent studies have investigated the predictability of adversarial developmental outcomes linked to prematurity. The purpose is to identify biomarkers in order to set up appropriate subject-specific interventions to counter or reduce the effects of neurodevelopmental afflictions. Several studies have focused on term-equivalent-age neuroimaging in order to evaluate the predictive capacity of either anatomical (Woodward et al., 2006), diffusion (Spittle et al., 2011) or functional imaging (Della Rosa et al., 2021) on motor and cognitive outcomes. A few studies have also investigated data acquired earlier than term-equivalent-age, and checked which has a better predictive power between early, term-equivalent, or the difference between them in case of longitudinal studies. Examples of such studies included motor outcome prediction, using volumetric measures relative to whole brain volumes (Gui et al., 2019), diffusion tensor parameters (Roze et al., 2015), or computing anatomical features such as cortical surface area or gyrification indices (Moeskops et al., 2017). All these studies focused on whole brain analysis to predict outcome. Yet, because of important methodology discrepancies, the results we obtained are hardly comparable.

We chose to investigate the *predictive ability of the central sulcus’ shape on motor development*, in the continuity of our shape characterization and description approach, in order to interrogate its functional relevance. In particular, we observed that the best shape feature to discriminate handedness was the length and backwards bend of the hand-knob of the left central sulcus at 30w PMA, and that a poor fine motor outcome at 5 years was mostly linked to a higher, deeper, but shorter right hand-knob, with the sulcal upper part bending forwards, at 40w PMA. To the best of our knowledge, we are the very first to assess the predictive capacity of sulcal shape during early development. We decided to use a classifier trained on the clinical factors that have been reported as relevant for motor outcome prediction in previous studies (Anderson et al., 2006; Kersbergen et al., 2016) as a benchmark to evaluate whether some shape features were relevant in classifying motor outcome, but other clinical factors might have been relevant to include, such as white matter injuries or venous infarctions. By extension we aimed to assess if motor neurodevelopmental adversities showed a signature in the central sulcus. In this “proof-of-concept” study, it seems that a few early sulcal features, in some hemisphere- and age-specific groups, contain functional information; yet, approximately half of the results reported showed a ROC AUC inferior to 0.5 (indicating a classifier performing worse than random). This can be due to an absence of relevant data to predict motor outcome in the shape of the central sulcus on specific groups, or to a methodological choice taken in this article: the predictors are probably under-fitted. Because of the size of the cohort and the small proportion of either left-handers or poor-manual-dexterity-outcomes, the decision was taken to prevent from operating feature selection or algorithmic tuning such as regularization in order to prevent from overfitting. We can also question the way we decided to dichotomize outcomes.

A limitation to our classification is that the data is globally preprocessed before entering the K-Fold cross-validation process (i.e. the Isomap dimensions are computed using the whole cohort before the classification process). This is discouraged as it induces bias in the classification. The recommended method would be to produce new Isomap dimensions at each fold, using only the training set. Yet, we chose to pursue the first method nonetheless, to enable interpretation: if we generated new Isomap dimensions at each fold, we would be unable to associate a given feature to each dimension, and dimensions would risk swapping within different folds. This would have prevented us from grasping the relevant shape features.

In terms of *handedness*, we observed that the left hemisphere was more informative than the right one, and the only relevant classifier using only shape features was obtained at 30w PMA, where left-handers tend towards a backwards bend of the superior part of the left central sulcus and a longer hand-knob (dimension 10). The result concerning the length of the hand-knob could suggest that, while the shape of the hand-knob is mostly driven by the hand area in the dominant hemisphere, the bulk of the hand-knob in the non-dominant hemisphere could be influenced by the adjacent functional areas (such as the immediate upper portion corresponding to the arm), justifying its wider sprawl. As indicated in the Methods section, for the analysis we considered as left-handers only the left-handed children who had at least one left-handed parent. Taking into account a genetic factor was an attempt to retain as much “natural” left-handers as possible (i.e. subjects who were most likely to present this handedness independently from the pathological implications of extreme prematurity). Interestingly, the number of left-handers obtained with this criterion corresponds to the range usually observed in the general population (7/71: 9.9%). Nevertheless, it has been reported that additive genetic effects only accounts for a quarter of the variance in handedness (Medland et al., 2009), therefore this left-hander-selection-choice is somewhat arbitrary. However, recent studies reported that paternal non-right-handedness was significantly related to left-hand preference in children (van Heerwaarde et al., 2020; Fagard et al., 2021). In our study, all the selected left-handers appeared to have a left-handed father, comforting our criteria choice. Still, there is no way to confirm that the 7 left-handers that we studied for the laterality analysis are “natural” left-handers. To look into this, we observed a condition which can alter lateralization towards left-handedness: left-hemisphere IVH. The excluded left-handers showed a higher prevalence of left-sided IVH than the selected left-handers (see Sup. Info., Table 5), suggesting that our criteria for selection was relevant to an extent. Yet, the selected left-handers showed a higher prevalence of left-sided IVH than the right-handers, as well as a globally worse MRI anomaly classification using the Kidokoro scale (Kidokoro et al., 2013) (see Sup. Info., Table 5). Therefore, we might not have succeeded in filtering adversarial left-handedness. This is confirmed by our observations, since the best classifiers for handedness were obtained by including clinical factors, which should have been irrelevant for “natural” handedness classification. In order to dim out pathological variability, further studies should focus on cohort selection, either by selecting preterms depending on clinical factors, or by looking into predictability of handedness through the central sulcus’s shape on term-born neonates.

In terms of *motor outcome prediction*, we observed relevant sulcal signatures at both ages. We aimed to investigate fine motor outcome (and not global motor score) in order to focus on the development of manual dexterity, as we were interested on the central sulcus shape that is highly characterized by the hand-knob, whose shape is associated with hand activation (Sun et al., 2016; Germann et al., 2019; Mangin et al., 2019). We should highlight that in the mABC-II, the subscores have not been thoroughly tested for psychometric relevance, contrarily to the global score (Hirata et al., 2018). They still have been reported to be acceptably relevant in different studies, especially for the manual dexterity and balance subscores (Ellinoudis et al., 2011). It is for all these reasons that we specifically chose to focus only on the manual dexterity subscore. The most weighted shape feature for fine motor outcome (dimension 10) was observed at 40w PMA on the right hemisphere, suggesting a narrower hand-knob corresponding to the left hand in infants with poor fine motor outcome. This could suggest a compensation for a less functional right hand, but this hypothesis needs to be investigated more thoroughly, especially since we did not select only right-handers for the fine motor outcome classification. Interestingly, this same 10^th^ dimension was the most weighted for the handedness on the left 30w PMA central sulcus (tendency to a longer hand-knob and an upper part of the sulcus bending backwards in left handers). This might suggest a less precise location of the right-hand region at 30w PMA. Although relevant for both motor classifications, this 10^th^ dimension shows no significant correlation between 30 and 40w PMA sulci: the early 30w shape encoded in the sulcus does not evolve to the corresponding shape at 40w PMA. A recent study refined the somatomotor mapping of the central sulcus and differentiated the central sulcus in 5 segments (Germann et al., 2019). Here, the upper part of the sulcus, bending forwards to backwards along the dimension, fits the first segment reported in that study, and the second segment corresponds to the hand area and comprises the hand-knob area. Segments one and two are reported to be separated by a narrow gyral passage linking the precentral and postcentral gyrus. It is possible that this buried gyrus develops mostly between 30 and 40w PMA and would justify the discordant expression of this feature at 30 and 40w PMA.

To sum up, this study, based on an original method for sulcal investigation, allowed us to describe the central sulcus’ shape variability and consistency at 30w and 40w PMA in a cohort of infants born extremely preterm and to make exploratory investigations of the functional relevance of these features. To complement our study, it would be relevant to compare the central sulcal shape of preterm infants at TEA with that of term-born newborns imaged right after birth, in order to assess whether prematurity-induced patterns are captured by our method. It is interesting to note that both for handedness and fine motor outcome, the left central sulcus at 30w PMA seemed to carry relevant information, which suggests that these functional aspects are already partially encoded at this early age. This opens perspectives about the possibility to target subjects at risk of abnormal motor development very early on and to mitigate this risk through early interventions.

## Acknowledgments

The authors thank Y. Leprince, H. Peyre, L. Perus, L. Guillon, C. Poiret and W. Shu-Quartier-Dit-Maire for their participation in discussions about this study.

## Supplementary Information

**Supplementary Table 1:** Perinatal clinical characteristics and fine motor follow-up of the study participants (n=71)

| *Characteristics* | *Mean ± Standard deviation (range) or N (percentage)* |
| --- | --- |
| *Perinatal clinical characteristics* |  |
| Sex, male | 36 (51%) |
| Gestational age at birth (weeks) | 26.5 ± 1.0 (24.4 – 27.9) |
| Birth-weight z-score | 0.4 ± 0.8 (-2.5 – 1.8) |
| Presence of severe IVH (grade 3 or 4): number of infants | 8 (11%) |
| Presence of broncho-pulmonary dysplasia: number of infants | 20 (39%) |
| *Age at MRI scans* |  |
| PMA at early acquisition (weeks) | 30.7 ± 0.9 (28.7 – 32.7) |
| PMA at term-equivalent age acquisition (weeks) | 41.2 ± 0.6 (40.0 – 42.7) |
| *Fine motor follow-up at 5-years* |  |
| Manual lateralization |  |
| Handedness: number of infants (left / ambidextrous / right) (n=70) | 18 / 2 / 50 (26 / 3 / 71%) |
| Corrected handedness*: number of infants (left / right) (n=57) | 7 / 50 (12 / 88%) |
| Fine motor assessment (n=66) |  |
| Age at mABC | 5 years 9 months ± 4 months (4y 6m – 6y 7m) |
| mABC manual dexterity standardized score | 7.7 ± 2.4 (3 – 14) |
| mABC manual dexterity outcome: number of infants (poor/borderline/good) | 15 / 16 / 35 (23 / 24 / 53%) |

IVH: intra-ventricular hemorrhage. PMA: post-menstrual age. \*Corrected handedness excludes ambidextrous and left-handed children having both parents right-handed. mABC: Movement Assessment Battery for Children.

**Supplementary Table 2:**
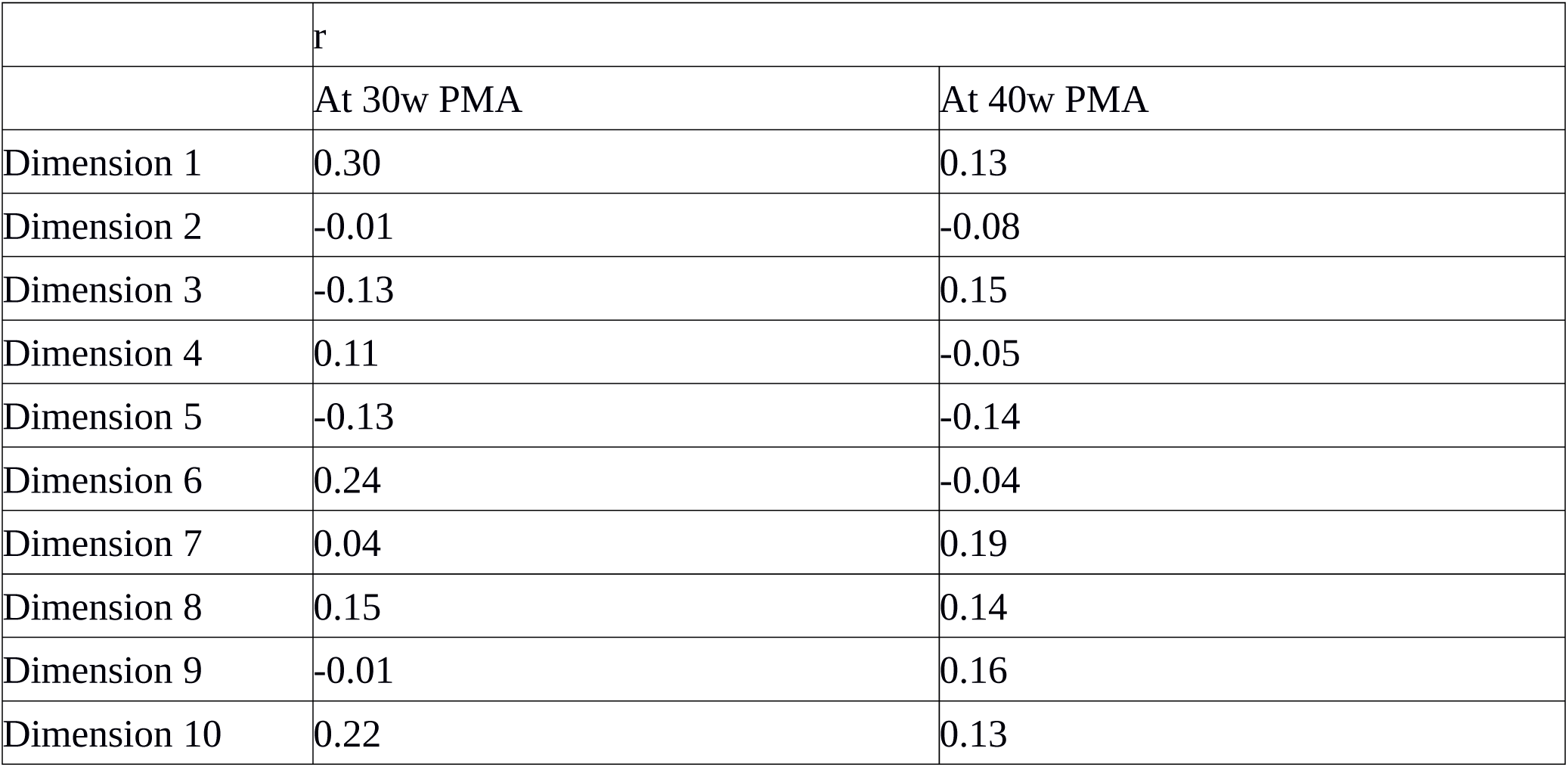
Pearson correlations between Isomap positions and PMA at MRI acquisition (n=71)

**Supplementary Table 3:** Wilcoxon signed-rank tests between either left / right hemisphere (A.) or 30w / 40w PMA (B.), for Isomap PMA-corrected positions (n=71)

|  | Wilcoxon signed-rank test: t (p-value) |  |  |  |
| --- | --- | --- | --- | --- |
|  | A. Hemispheric comparison |  | B. Age-group comparison |  |
|  | L30 vs R30 | L40 vs R40 | L30 vs L40 | R30 vs R40 |
| Dimension 1 | 959 (0.068) | 1022 (0.142) | **3.0 (3.10<sup>-13</sup>)** | **0.0 (2.10<sup>-13</sup>)** |
| Dimension 2 | 1044 (0.180) | 1276 (0.991) | 882.0 (0.023) | **774.0 (0.004)** |
| Dimension 3 | **757 (0.003)** | 1110 (0.336) | 903.0 (0.032) | 978.0 (0.086) |
| Dimension 4 | 941 (0.053) | 1113 (0.344) | **627.0 (2.10<sup>-4</sup>)** | 1129.0 (0.393) |
| Dimension 5 | 878 (0.022) | 1137 (0.419) | **606.0 (1.10<sup>-4</sup>)** | **686.0 (7.10<sup>-4</sup>)** |
| Dimension 6 | 1133 (0.406) | 1152 (0.470) | 1165.0 (0.517) | 1178.0 (0.567) |
| Dimension 7 | 1169 (0.532) | 947 (0.058) | 986.0 (0.094) | 849.0 (0.014) |
| Dimension 8 | 1242 (0.837) | **775 (0.004)** | 804.0 (0.007) | 1138.0 (0.422) |
| Dimension 9 | 1208 (0.689) | 1254 (0.891) | 1099.0 (0.305) | 1212.0 (0.705) |
| Dimension 10 | 1132 (0.403) | 881 (0.023) | 871.0 (0.020) | 1066.0 (0.224) |

Bold values verify p-value<0.005. L: left hemisphere; R: right hemisphere. 30: 30w PMA (early scan); 40: 40w PMA (term-equivalent-age scan).

**Supplementary Table 4:** Spearman correlations between either left / right hemisphere (A.) or 30w / 40w PMA (B.), for Isomap PMA-corrected positions (n=71).

|  | Spearman correlation: ρ (p-value) |  |  |  |
| --- | --- | --- | --- | --- |
|  | A. Hemispheric comparison |  | B. Age-group comparison |  |
|  | L30 vs R30 | L40 vs R40 | L30 vs L40 | R30 vs R40 |
| Dimension 1 | **0.607 (2.10<sup>-8</sup>)** | **0.385 (9.10<sup>-4</sup>)** | 0.232 (0.051) | **0.428 (2.10<sup>-4</sup>)** |
| Dimension 2 | **0.407 (4.10<sup>-4</sup>)** | 0.232 (0.051) | **0.458 (6.10<sup>-5</sup>)** | 0.329 (0.005) |
| Dimension 3 | 0.222 (0.063) | **0.351 (0.003)** | 0.307 (0.009) | **0.353 (0.003)** |
| Dimension 4 | 0.254 (0.032) | 0.283 (0.017) | **0.512 (5.10<sup>-6</sup>)** | **0.414 (3.10<sup>-4</sup>)** |
| Dimension 5 | **0.411 (3.10<sup>-4</sup>)** | 0.327 (0.005) | **0.376 (0.001)** | 0.317 (0.007) |
| Dimension 6 | 0.216 (0.070) | 0.150 (0.213) | 0.284 (0.017) | **0.383 (0.001)** |
| Dimension 7 | 0.188 (0.116) | 0.249 (0.036) | 0.081 (0.500) | 0.191 (0.111) |
| Dimension 8 | 0.212 (0.076) | 0.186 (0.120) | 0.246 (0.039) | **0.522 (3.10<sup>-6</sup>)** |
| Dimension 9 | 0.157 (0.190) | 0.123 (0.308) | 0.294 (0.013) | **0.334 (0.004)** |
| Dimension 10 | 0.158 (0.189) | 0.264 (0.026) | -0.024 (0.842) | -0.063 (0.599) |

Bold values verify p-value<0.005. L: left hemisphere; R: right hemisphere. 30: 30w PMA (early scan); 40: 40w PMA (term-equivalent-age scan).

**Supplementary Table 5:** Additional perinatal characteristics reported for right-handers, selected left-handers and excluded left-handers (based on parental handedness) (n=68)

| Characteristics | N (percentage) |  |  |
| --- | --- | --- | --- |
| Handedness class | Right-handers (n=50) | Selected left-handers (n=7) | Excluded left-handers (n=11) |
| Presence of left IVH (any grade): number of infants | 12 (24%) | 2 (29%) | 4 (36%) |
| Kidokoro classes |  |  |  |
| Normal | 25 (50%) | 1 (14%) | 3 (27%) |
| Mild | 20 (40%) | 6 (86%) | 7 (64%) |
| Moderate | 4 (8%) | 0 (0%) | 1 (9%) |
| Severe | 1 (2%) | 0 (0%) | 0 (0%) |

IVH: intra-ventricular hemorrhage

## Annex 1 : Parameter selection for the Isomap algorithm

### Method to choose the intrinsic dimensionality of the manifold *dopt* for a given *k*

Let M*_dist_*be the input distance matrix. A distance matrix of D-dimensional Gaussian vectors was computed, with the same average square distance as *M_dist_*, which we called *M_rand._* Using a loop, we computed the reconstruction error *e_dist_*of the Isomap fitted on *M_dist_* and the reconstruction error *e_rand_* of the Isomap fitted on *M_rand_* for the whole range of possible number of components for the Isomap, so for d∈[1,*dim*(*M_dist_*)]. We then considered the number of components maximizing the ratio *e_rand_/e_dist_*as the intrinsic dimensionality of the manifold (see Supplementary Figure 1.1.A).

### Method to choose the number of nearest neighbors *k*

Using the previous method, we computed the intrinsic dimensionality of the manifold for each possible *k* in our study, so for *k*∈[1, 283]. The choice of *k* was based on the following considerations:

1. In order to prevent short-circuits in the manifold, *k* should be rather small.
2. The smallest value to consider for *k* is the smallest value allowing the nearest neighbors graph to be connected ⇒ *k*min = 2.
3. Such a small *k* is not suited for our study since we wish to interconnect 30w sulci with 40w sulci and reciprocally. The number of inter-age-subgroup connections grows with *k* (see Supplementary Figure 1.1.B).

**Supplementary Figure 1.1.**
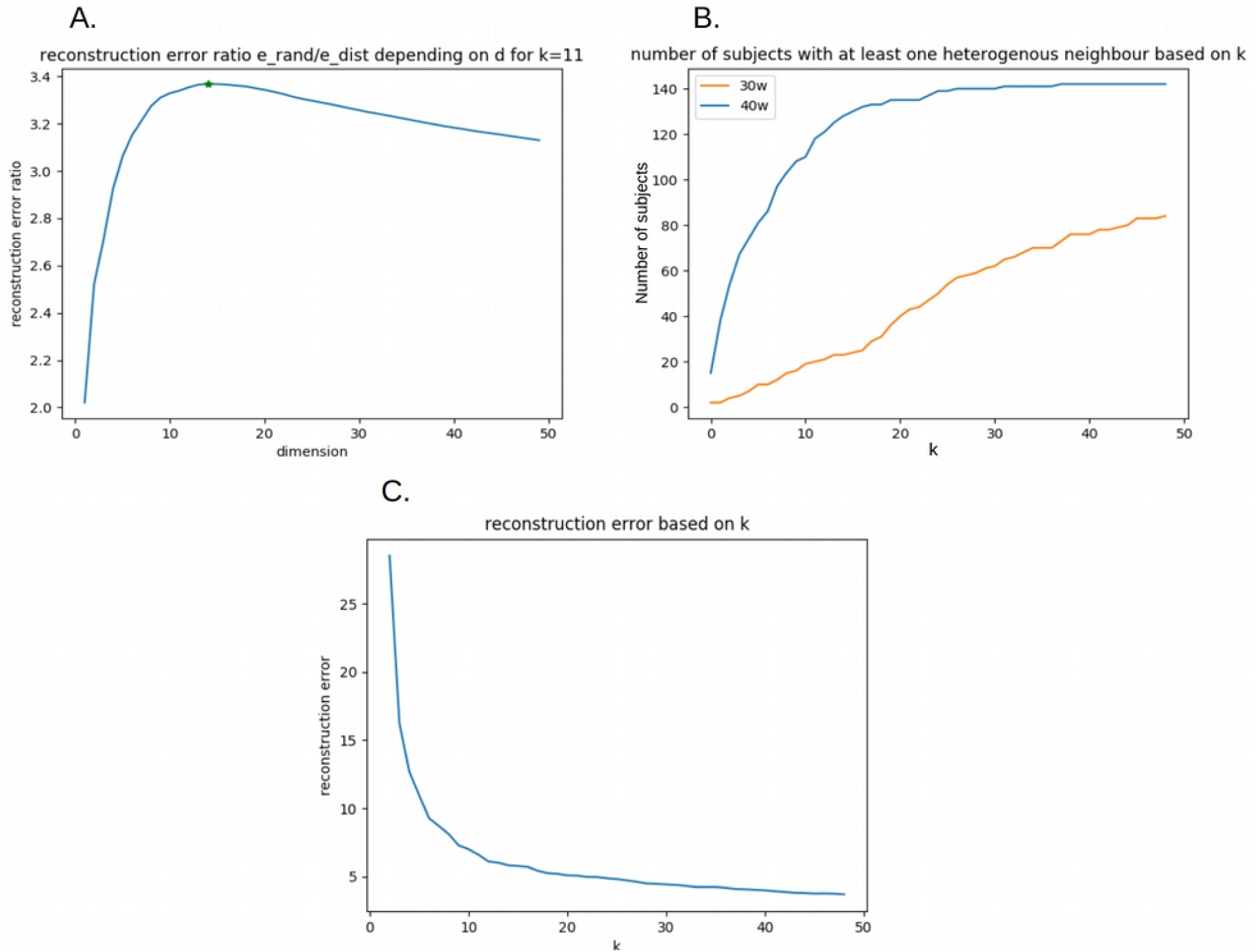
A. Reconstruction error ratio depending on dimension *d* for a fixed *k* (here, *k*=11, leading to an intrinsic dimensionality of 14). B. Number of sulci with at least one nearest neighbor of the other age-group depending on *k*. Orange: 30w sulci, blue: 40w sulci. C. Reconstruction error based on *k*.
4. Our purpose was to choose a *k* small enough to prevent short-circuits in the manifold, but high enough to capture accurately the complexity of the dataset. In order to get an idea of the range of *k* which corresponds to this criterion, we have plotted the graph of reconstruction error depending on *k* (see observed that the Isomap results Supplementary Figure 1.1.C). We observed an interesting reconstruction error drop between *k*=5 and *k*=20, so this is the scope we focused on.
5. In this scope, we have observed that the Isomap results did not differ drastically, as can be seen using the correlation matrices for the Isomap projections using different values for *k*, as shown in Supplementary Figure 1.2.

**Supplementary Figure 1.2.**
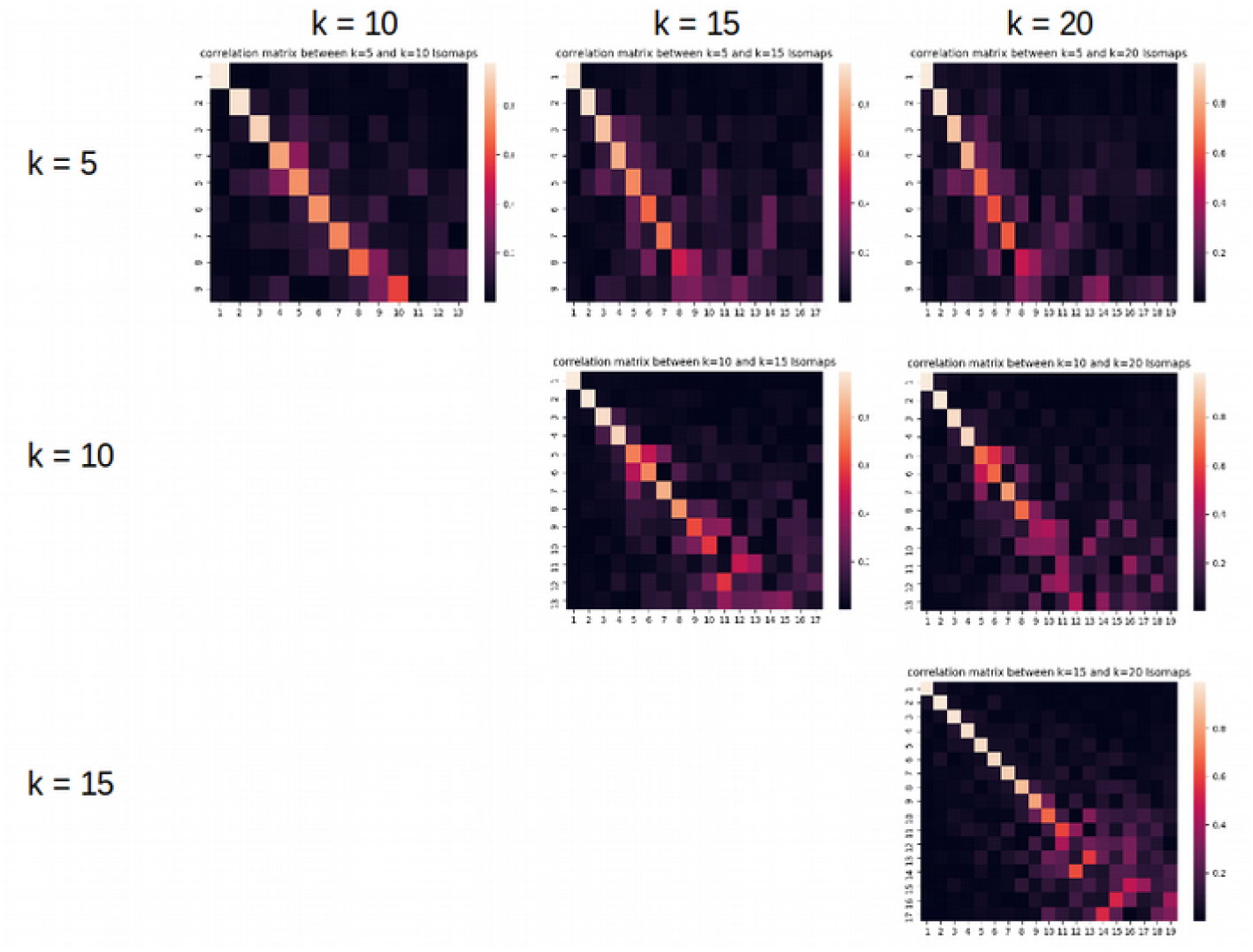
Correlation matrices between the positioning of the sulci on the dimensions obtained using *k*∈{5,10,15,20}. The correlation values suggest that most dimensions are not drastically altered by different choices of *k* in this scope.
6. In order to select a specific *k* within this range, we looked into the residual variance graph based on k, as shown in Supplementary Figure 1.3.

**Supplementary Figure 1.3.**
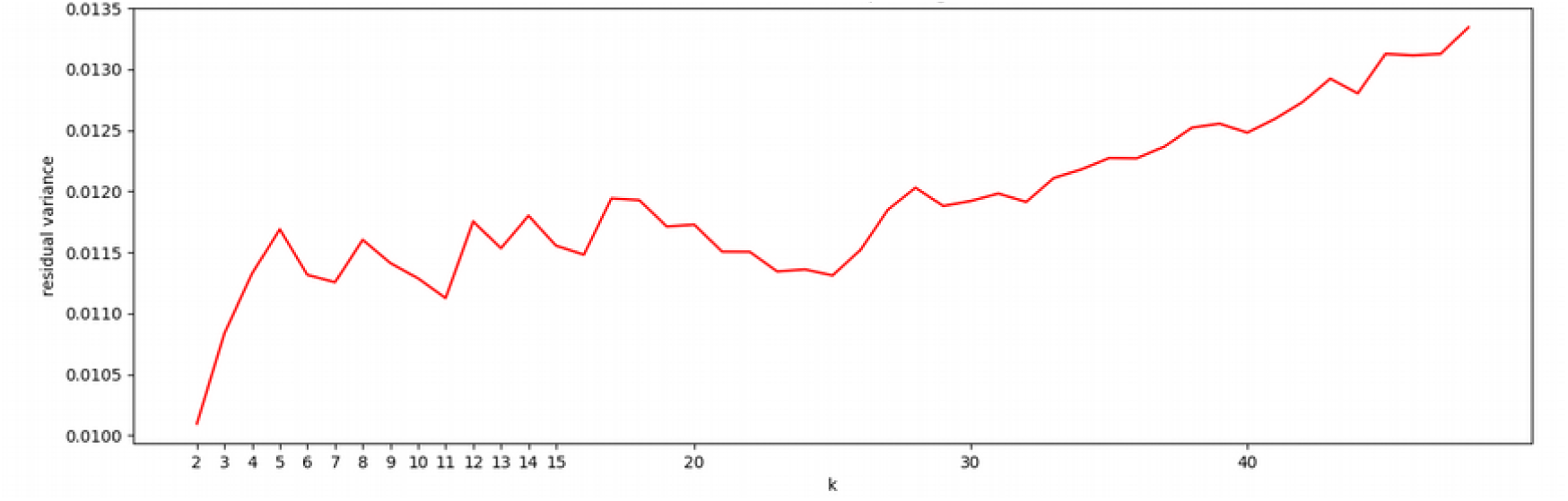
Residual variance depending on k.

We observed a local minimum for *k*=11 and therefore chose this value.

### Method to choose the relevant number of dimensions for our study

According to Supplementary Figure 1.1.A, the intrinsic dimensionality for *k*=11 was 14. Looking back into this figure, we observed that the reconstruction error ratio reached a plateau. The fact that *e_rand_/e_dist_* did not change around this plateau suggested that the additional data gained on the input distance matrix was not significantly higher than the random distance matrix. Therefore, in order to capture the most relevant dimensionality reduction in the data, we considered that even though the intrinsic dimensionality of the manifold was the one maximizing this ratio, the relevant dimensionality to consider in this study was the first *d* for which the curve stopped growing significantly. Here, by looking at the derivative from figure Supplementary Figure 1.4, we considered the relevant dimensionality *d*=10:

**Supplementary Figure 1.4.**
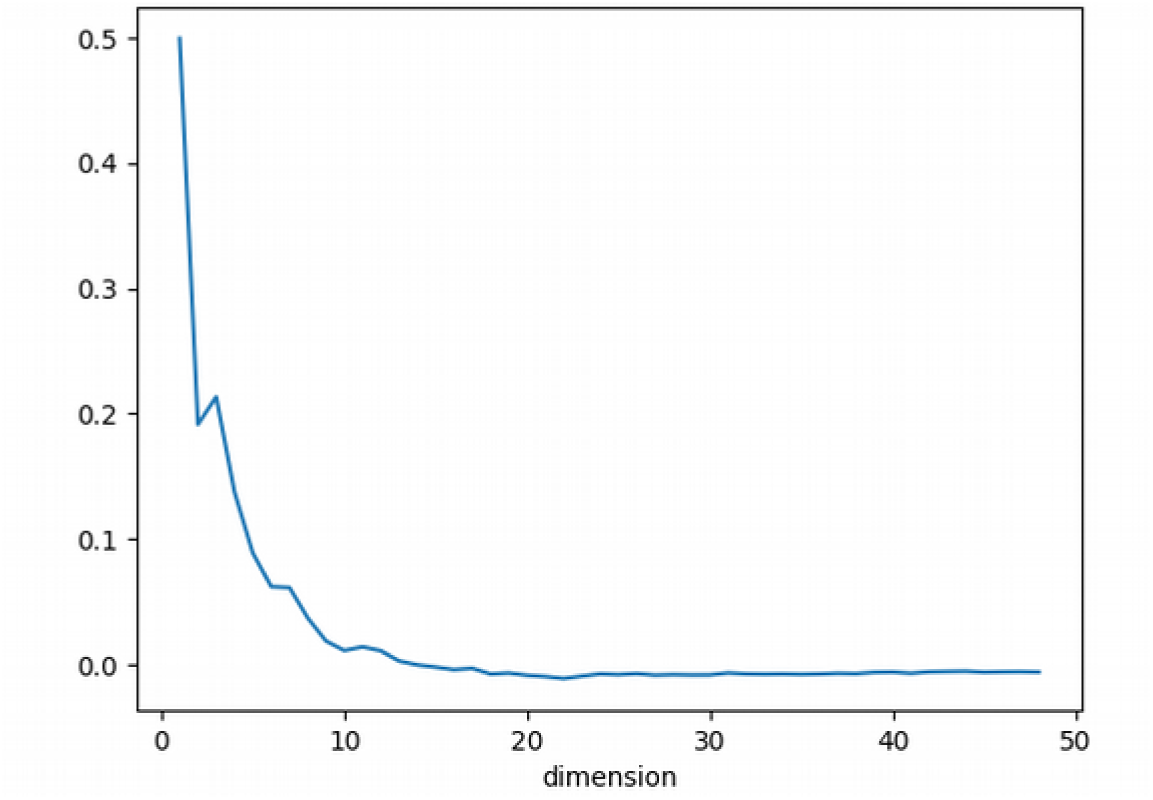
Derivative of the reconstruction error ratio for *k*=11 depending on *d*

### Confirmation that the observations for *k*=11 (and *d*=10) are stable for neighboring values of *k*

To confirm that this decision is not inducing too much bias in our observations, we computed the correlation matrices for the Isomap projections using *k*=10 and *k*=12 to compare with *k*=11, represented on Supplementary Figure 1.5.

**Supplementary Figure 1.5.**
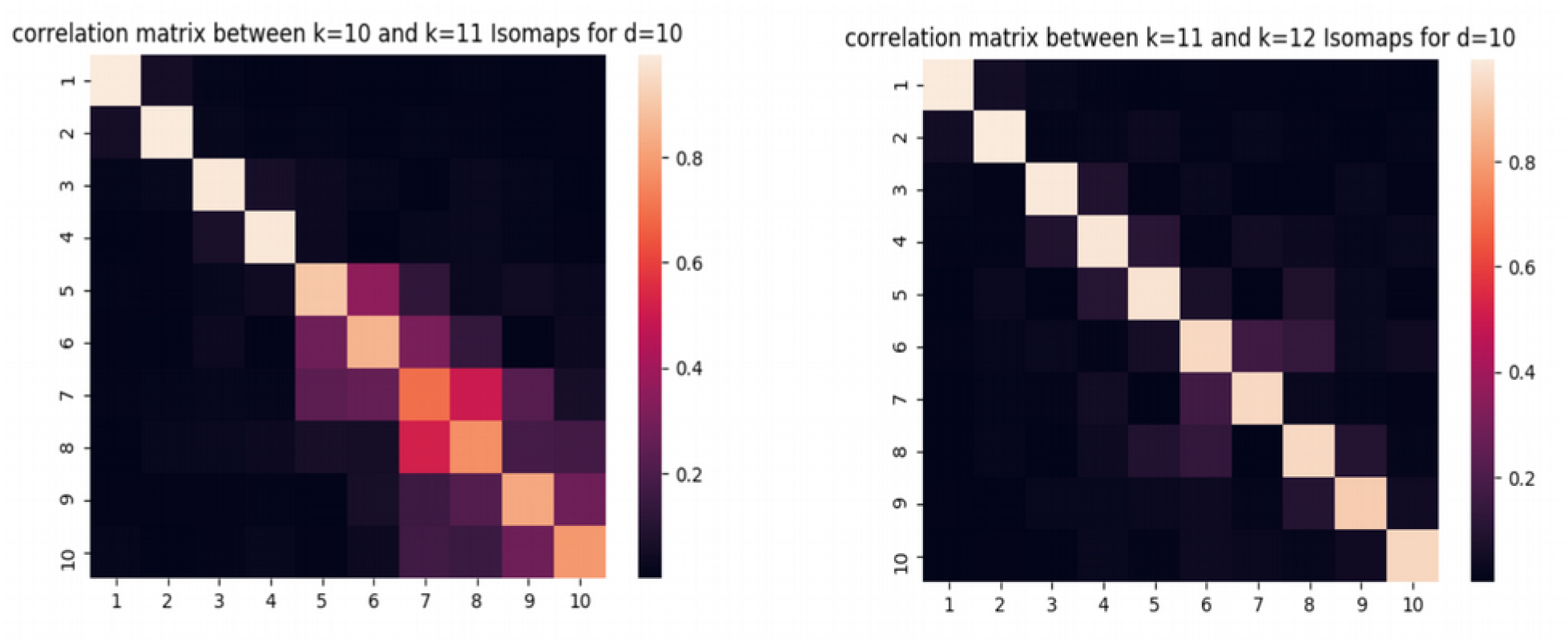
Correlation matrices between the positioning of the sulci on the dimensions obtained using *k*=10 and *k*=11 (left), and *k*=11 and *k*=12 (right).

These correlations matrices confirmed that the choice of parameter *k*=11 does not excessively affect the projection results.

## Annex 2 : Representation of the 10 Isomap dimensions

For each dimension, the sulcal projections were represented as well as the age-specific moving averages (blue = 30w PMA, green = 40w PMA). A description for the shape feature captured is suggested for each dimension. Additionally, the information about statistical tests for each dimension (Wilcoxon signed-rank test and Spearman correlation for L30, R30, L40 and R40) is summarized in the table next to it (✓for a p-value < 0.005, ∼ for a p-value between 0.05 and 0.005, and x for a p-value > 0.05).

The reading key for shape description (Figure 3.A) is reminded on Supplementary Figure 2.1.

**Supplementary Figure 2.1.**
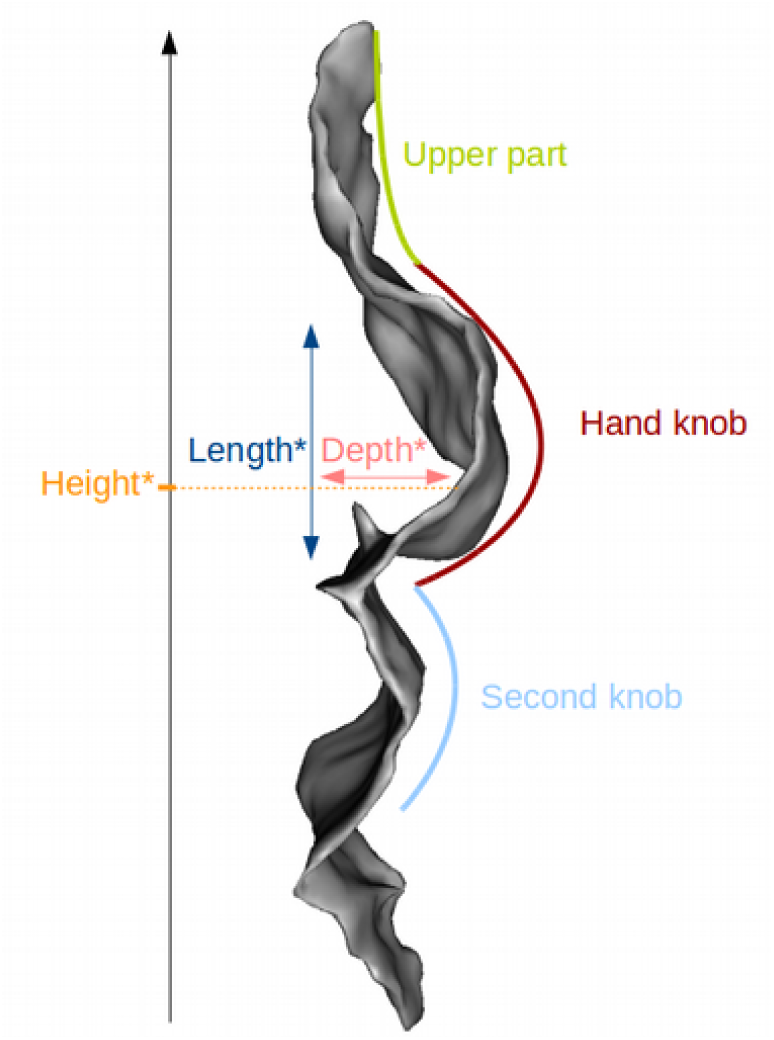
Reading key for shape description of the central sulcus with a two-knob configuration. The height of the hand-knob is defined as the vertical distance from the bottom of the sulcus to the deepest region of the hand-knob. *Measures relative to the hand-knob.

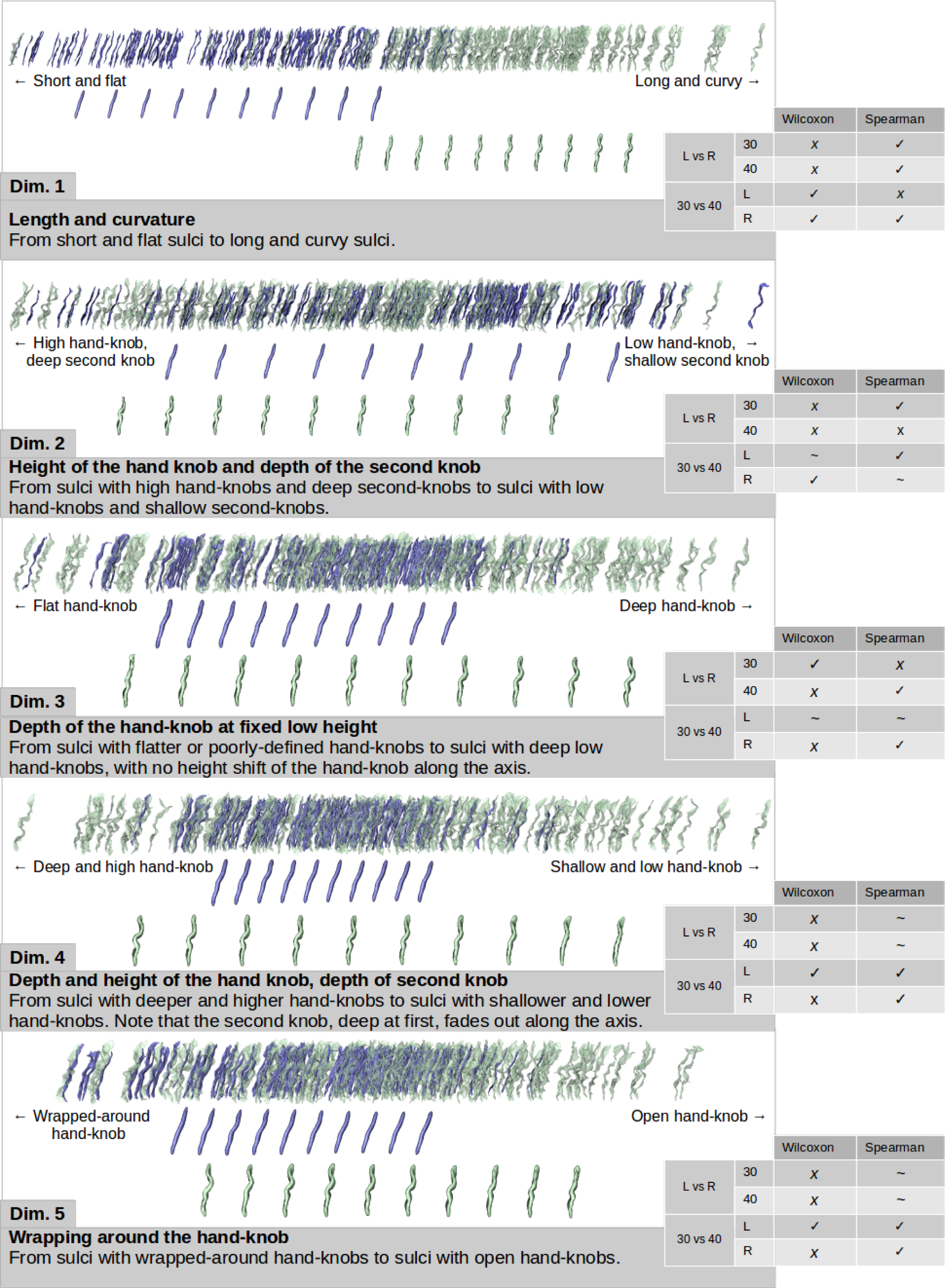
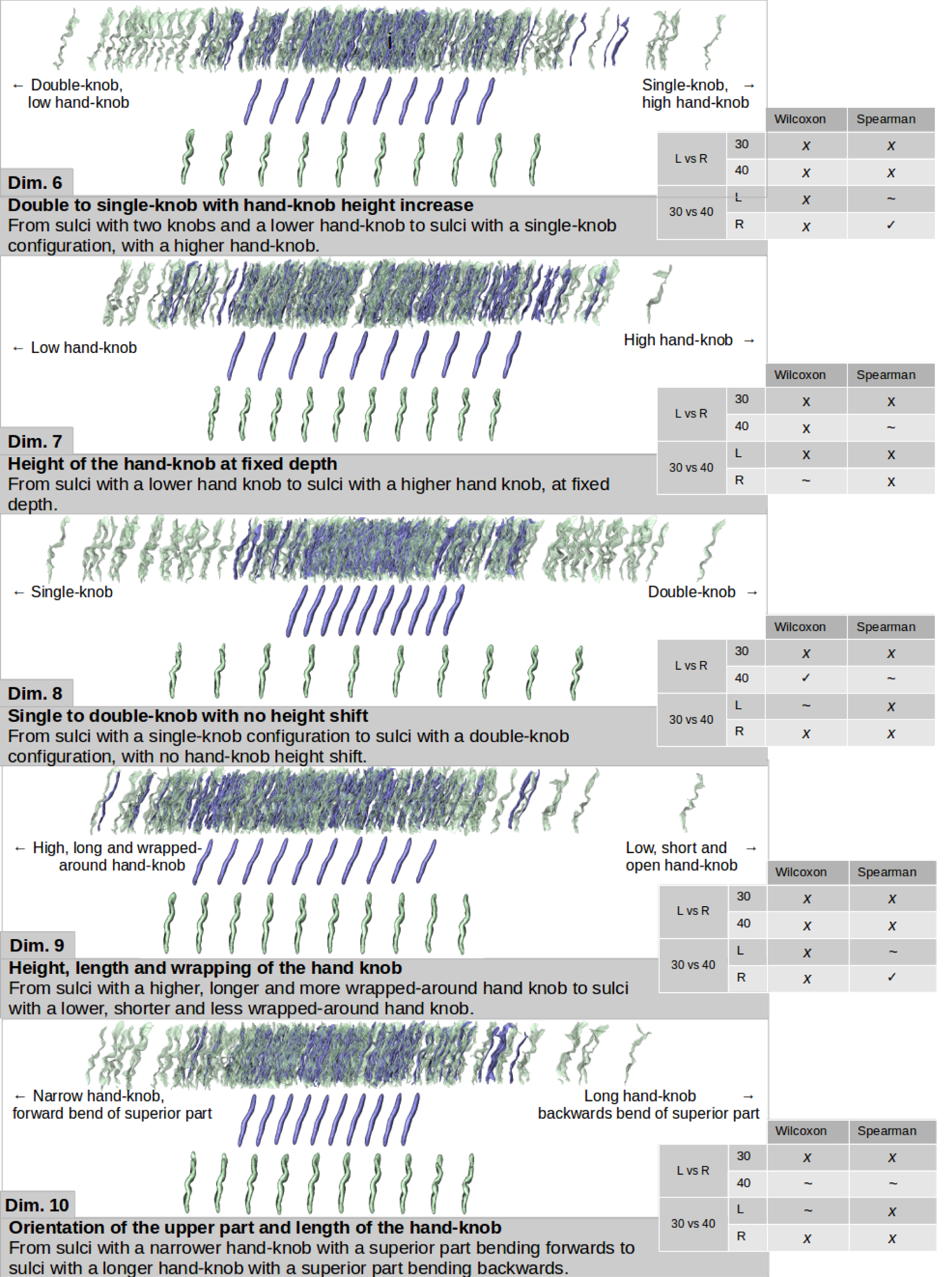

## Annex 3 : Analysis of the second-best mABC fine motor outcome classifier obtained at 30w PMA on the left hemisphere using sulcal shape features alone

Two classifiers using sulcal shape features alone scored a tie in terms of ROC AUC for fine motor outcome classification. The one with the highest recall (right hemisphere, 40w PMA) was presented in the main section. Here is presented the analysis of the other classifier, obained at 30w PMA on the left hemisphere.

In order to address the best-scoring classifiers for fine motor outcome, the comparison for ROC and PR curves between the baseline classifier and the left hemisphere 30w PMA sulcal shape features’ classifier is presented here.

**Supplementary Figure 3.1.**
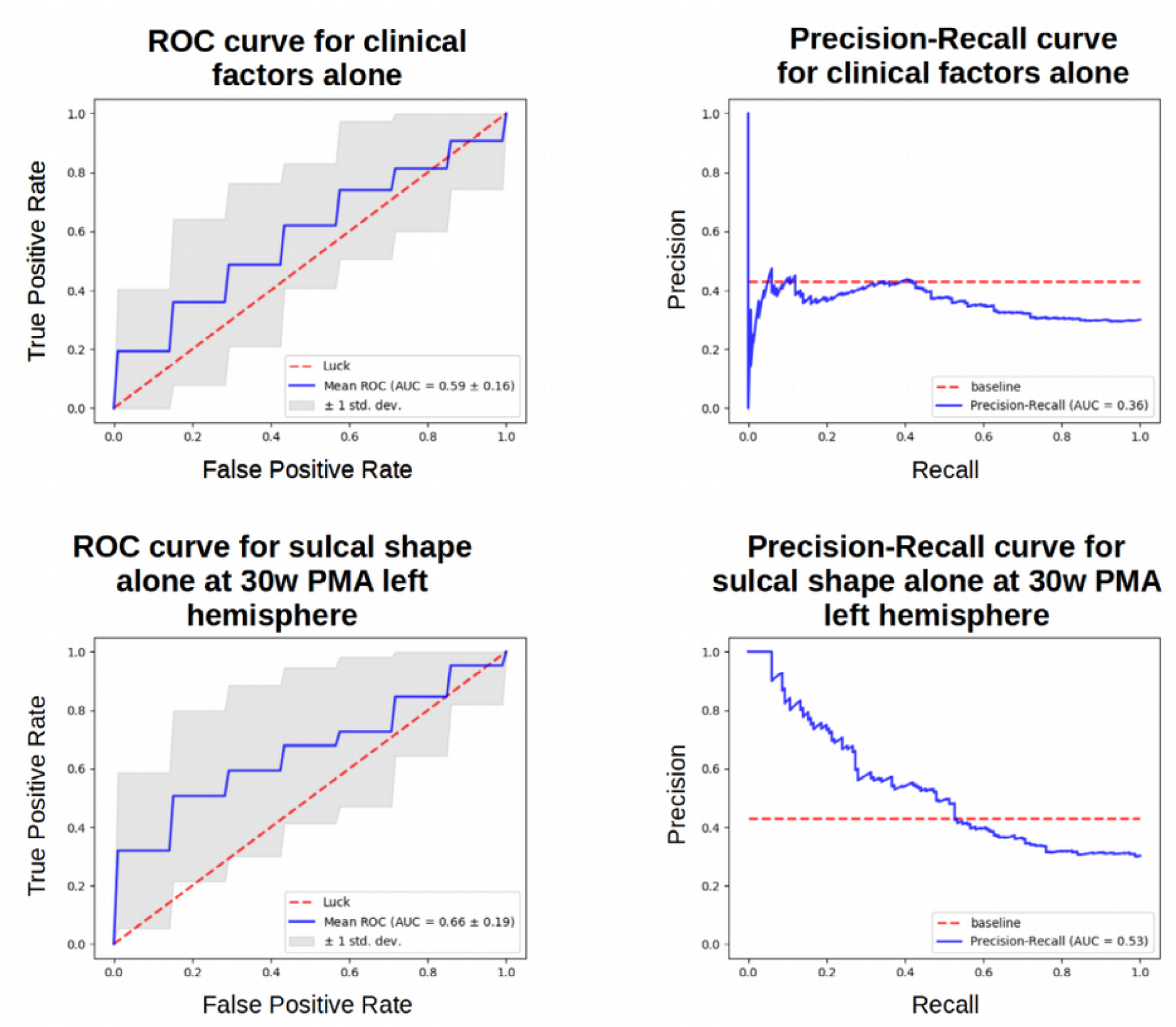
ROC and PR curves obtained for clinical factors alone and sulcal shape factors alone at 30w PMA on the left hemisphere. Target: fine motor outcome

Using the different folds of the repeated stratified cross-correlation, we aggregated the coefficients attributed to each feature and generated the box-plot (Supplementary Figure 3.2).

**Supplementary Figure 3.2.**
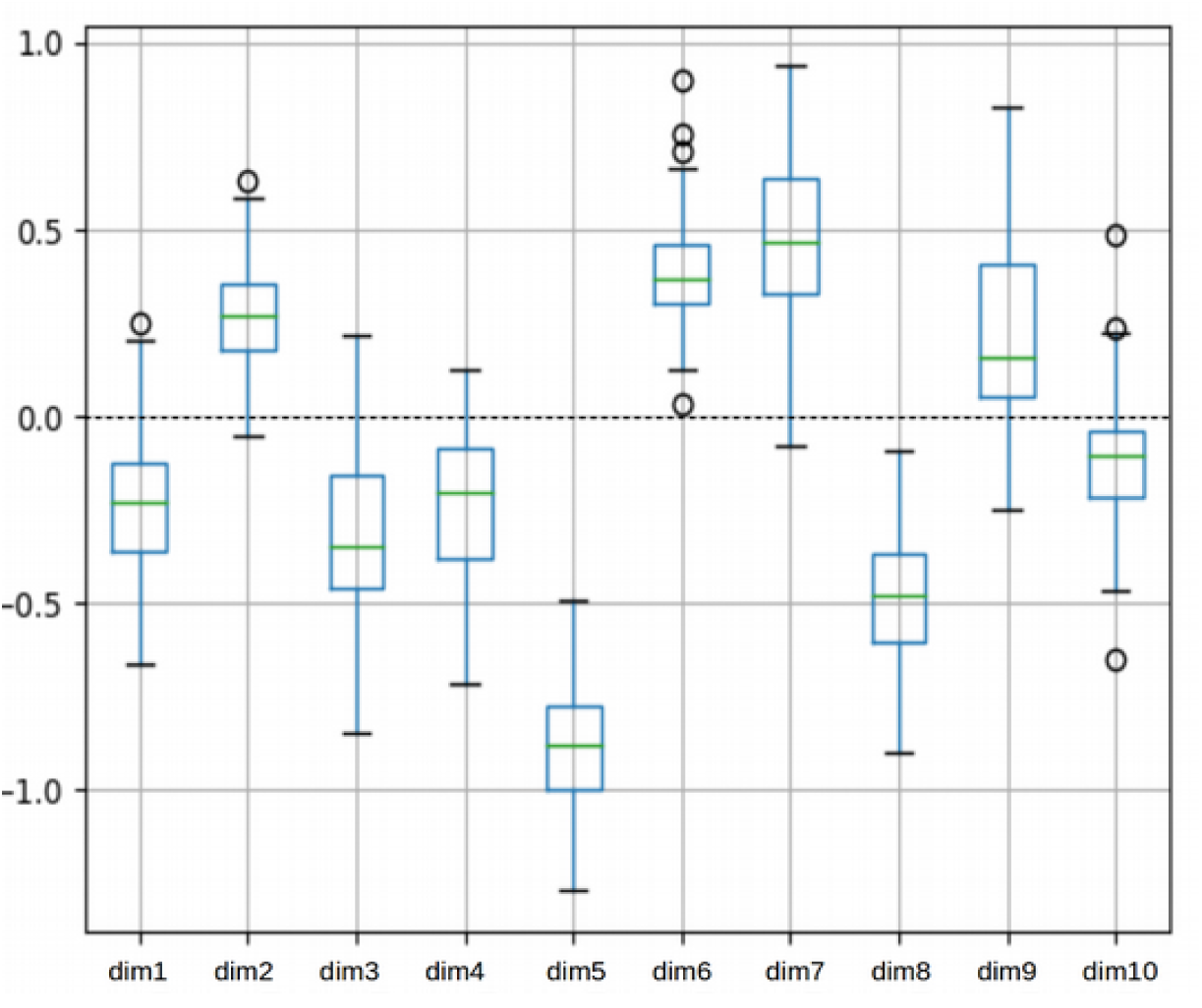
Box plot of the coefficients attributed to each Isomap dimension during the cross validation for the fine motor outcome classification with sulcal shape factors alone at 30w PMA on the left hemisphere. Coefficients for the dimensions were obtained for each iteration of a 5-fold 10-times repeated stratified cross-validation training of the SVC for fine motor outcome. Dimension 5 weighted generally more than the other dimensions throughout this cross-validation.

The most weighted feature appeared to be dimension 5, represented in Supplementary Figure 3.3. Compared to right-handers, the 30w PMA left central sulcus of left-handers tended to have a more wrapped-around hand-knob.

**Supplementary Figure 3.3.**
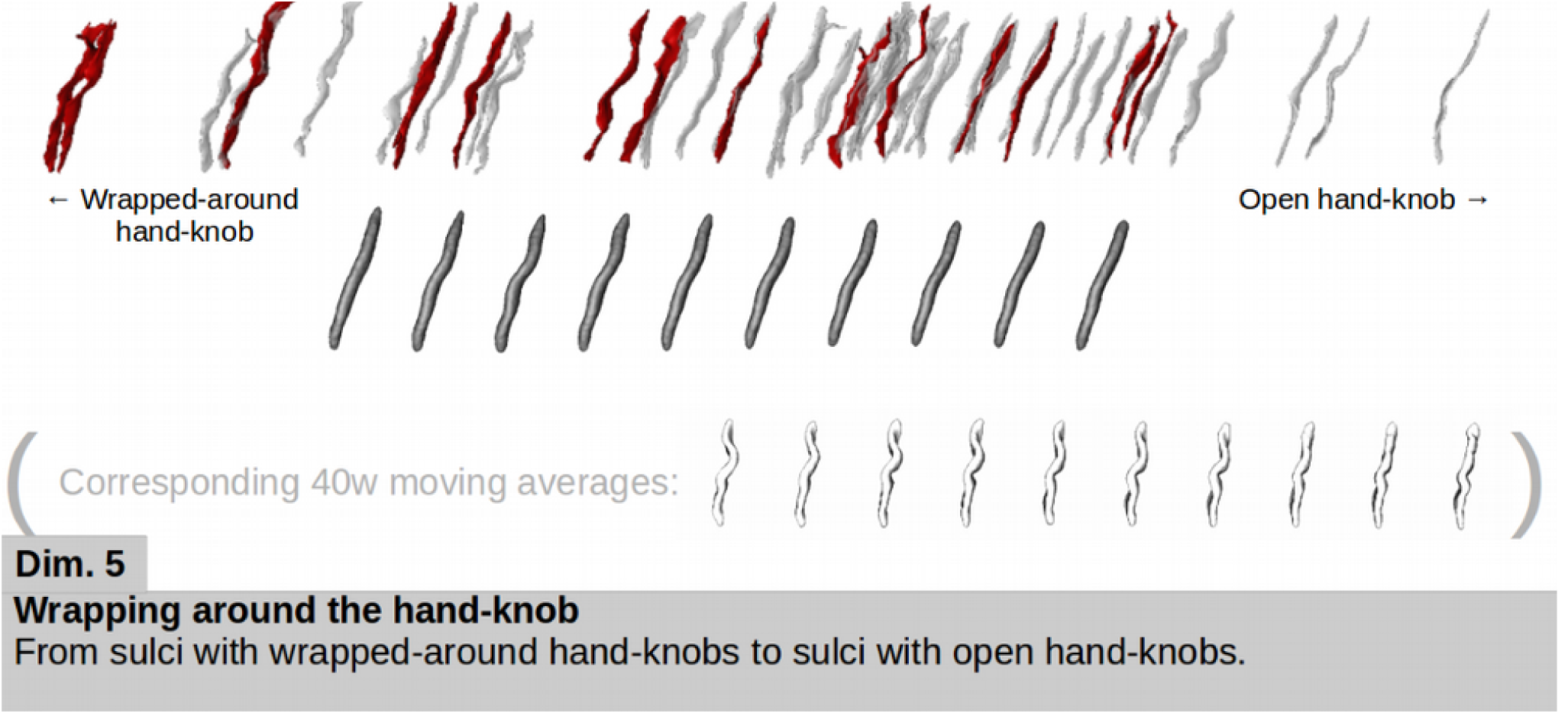
Representation of the dimension weighting the most for fine motor outcome classification using sulcal features at 30w PMA in the left hemisphere. The sulcal projections of the subjects used in the classification were represented (good outcome in grey, poor outcome in red), as well as the age-specific moving averages (in dark grey). The 40w PMA moving averages (in white) are shown for shape interpretation purposes.

